# NEXT-FRET: A solution-based smFRET platform to resolve folding intermediates under native conditions

**DOI:** 10.1101/2025.07.30.666321

**Authors:** Chara Sarafoglou, Andreas Kofidis, Marijn de Boer, Mikis Mylonakis, Kostas Mavrakis, Giannis Zacharakis, Yannis Pantazis, Giorgos Gouridis

**Affiliations:** Laboratory of Dynamic Structural Biology, Structural Biology and Biophysics Division, Institute of Molecular Biology and Biotechnology (IMBB-FORTH); Heraklion-Crete, 70013, Greece; Department of Biology, University of Crete; Heraklion-Crete, 70013, Greece; Institute of Applied and Computational Mathematics, Foundation for Research and Technology - Hellas; Heraklion-Crete, 70013, Greece; Department of Computer Science, University of Crete; Heraklion-Crete, 70013, Greece; Zernike Institute for Advanced Materials, University of Groningen, Groningen 9747 AG, The Netherlands; Laboratory for Biophotonics and molecular imaging, Institute of Electronic Structure and Lasers (IESL-FORTH), Heraklion-Crete, 70013, Greece; Kymatonics P.C., Heraklion-Crete, 70013, Greece

## Abstract

Folding intermediates are promising therapeutic targets in protein misfolding, bacterial virulence, and drug discovery. Yet, directly observing these transient, non-equilibrium states under physiological conditions remains challenging. Here, we introduce NEXT-FRET (Non-Equilibrium miXTure modeling of smFRET), a solution-based single-molecule FRET (smFRET) platform integrating accessible instrumentation and a time-variant Gaussian Mixture Model (tvGMM) to identify folding intermediates without microfluidics, surface tethering, or denaturants. Using maltose-binding protein (MBP) and its biologically relevant precursor (pre-MBP) for benchmarking and validation, we uncover previously elusive intermediates, including a long-hypothesized closed conformation undetected by existing single-molecule and ensemble methods. Furthermore, we demonstrate that chaperones modulate folding landscapes by stabilizing distinct intermediates, some effectively arresting folding progression; a mechanism increasingly recognized as a therapeutic strategy for targeted protein inactivation. Broadly applicable beyond folding, NEXT-FRET provides a generalizable, high-resolution analytical method for resolving transient biomolecular intermediates, offering a screening-compatible platform in biotechnology and elucidating proteostasis collapse in conformational diseases.

## Main

Understanding conformational dynamics in proteins is crucial to fundamental biology and translational biotechnology, especially where dynamics underpin folding, misfolding, disease etiology, and biological function. Single-molecule Förster Resonance Energy Transfer (smFRET) has become indispensable for directly probing conformational heterogeneity inaccessible to ensemble-averaged techniques^1^. A recent international blind study across 19 laboratories confirmed smFRET’s exceptional precision (≤2 Å) in quantifying protein conformational changes, cementing its role in structural biology^2^.

Biological processes predominantly occur far from thermodynamic equilibrium, involving transient intermediates and complex kinetics elusive to traditional methods. Protein folding exemplifies this complexity, where intermediate states dictate folding efficiency and fidelity^3,4^. Misfolding and aggregation can impair cellular homeostasis and contribute to neurodegenerative diseases^5,6^, underscoring the need for precise resolution of folding pathways. Yet, conventional ensemble techniques like NMR, SAXS, and crystallography while powerful for structural insights, obscure transient intermediates due to population averaging.

smFRET has revolutionized folding studies by capturing individual molecular folding trajectories in real time^7–10^. However, most implementations use surface-immobilization via Total Internal Reflection Fluorescence (TIRF) microscopy, introducing artifacts like surface-induced sticking that distort folding trajectories^11,12^ and limit observation times due to photobleaching; making it difficult to capture slow or rare folding events over extended periods^3,13^. Such artifacts can stabilize non-native conformations or restrict motion, reducing data accuracy and physiological relevance. While microfluidic mixing enables study of freely diffusing proteins at defined time points with millisecond temporal resolution ^1,7–10,14^; two key limitations constrain their broader utility. First, the fabrication of devises with sub-micron geometries demands specialized microengineering, limiting their accessibility to only highly specialized laboratories^15^. Second, the hydrodynamic conditions used to drive mixing introduces mechanical stress, not typical of physiological environments^16^, and may distort protein folding trajectories and/or chaperone association. Likewise, detergents are often used to stabilize unfolded or partially folded proteins and suppress aggregation, but they risk altering the folding landscape by promoting non-native interactions^17^. These challenges have hindered the broad application of smFRET to translational biology. To address these challenges, we introduce NEXT-FRET (Non-Equilibrium miXTure modeling of smFRET), a solution-based smFRET platform that circumvents surface tethering, microfluidics, and detergents. NEXT-FRET integrates minimally perturbing solution-phase measurements with a custom time-varying Gaussian Mixture Model (tvGMM), enabling resolution of transient folding intermediates and their temporal evolution without separating molecules in space or relying on artificial perturbants.

A central question in folding remains: do proteins fold through numerous independent pathways, as theoretical models suggest^18,19^, or via essential intermediates along a defined trajectory^20^, as experimental data suggest^21,22^? This distinction directly impacts drug discovery; as folding proceeds, essential intermediates represent kinetic bottlenecks and functional checkpoints, offering unique pharmacological intervention points^23^. Stabilizing these metastable states, particularly in virulence factors, disease-causing protein derivatives or aggregation-prone proteins, can inhibit (mal)function without requiring active-site binding^24^. Moreover, decoding the full spectrum of folding intermediates and their modulation by chaperones can reveal the molecular mechanisms underlying proteostasis collapse, central to many conformational diseases^25^.

Using maltose-binding protein (MBP)—a discontinuous two-domain protein extensively studied in folding research—we reveal a broader set of intermediates than previously detected. These states reconcile prior discrepancies and align with MBP’s free-energy landscape. Notably, NEXT-FRET remains robust in the presence of molecular chaperones and effectively resolves folding dynamics in pre-MBP, an aggregation-prone, signal-peptide bearing precursor previously inaccessible to single-molecule analysis. The simplicity, gentleness and resolution of NEXT-FRET thus provide a broadly generalizable platform for dissecting folding landscapes across diverse proteins relevant to biomedical and biotechnological application.

## Results

### Resolving non-equilibrium protein folding pathways using solution-based smFRET

#### smFRET experimental design and equilibrium benchmarks

To investigate MBP’s folding dynamics using smFRET, we employed site-specific labeling with donor and acceptor fluorophores. Fluorophores were positioned at the N- and C-termini to maximize distance resolution to large-scale folding transitions, and within individual domains to monitor conformational changes between the native open and closed states (Fig. 1a).

**Fig. 1.**
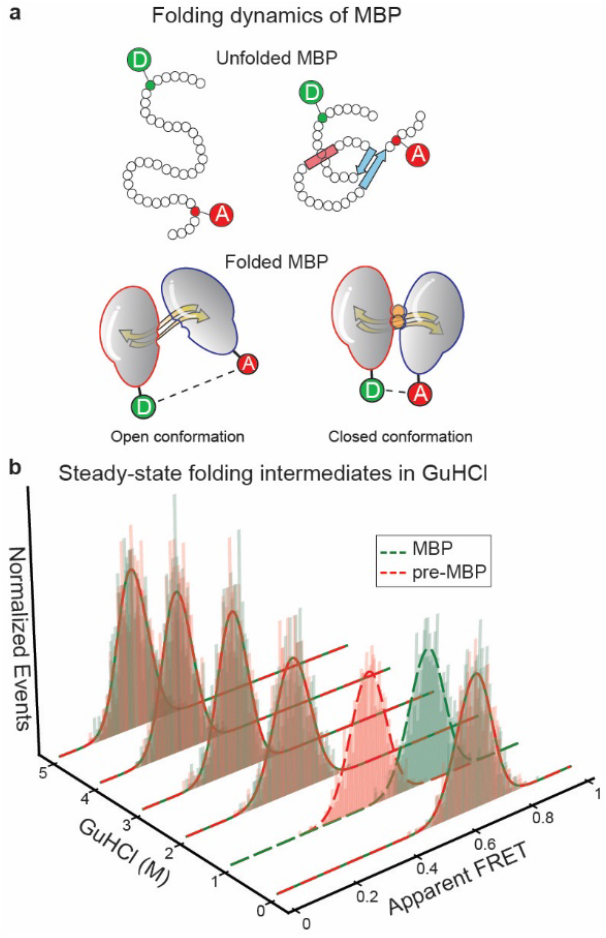
Experimental pipeline for establishing non-equilibrium smFRET analysis on folding. **a**, Donor (D) and acceptor (A) fluorophores are positioned at the N- and C-termini of the polypeptide chain to monitor intermediates during folding, with higher FRET values indicating closer proximity. Their strategic positioning also allows detection of native transitions from the open apo to the closed liganded conformation. **b**, Steady-state smFRET titrations with the chaotropic agent GuHCl used to define folding and unfolding conditions and confirm the recovery of the native state upon GuHCl concentration reduction, for MBP (green) and pre-MBP (red). Each Gaussian distribution represents the predominant conformational state at a given GuHCl concentration.

We first conducted two equilibrium control experiments to validate the functionality of the labeled proteins and to establish prior knowledge for NEXT-FRET. In the first, MBP was natively purified, labeled and measured under both apo-open (unliganded) and holo-closed (maltose-bound) conditions. These measurements confirmed that the labeled protein retained its structural and functional integrity and the ability to undergo maltose-induced conformational transition, in agreement with previous studies^26,27^. In the second control, we characterized the unfolding behavior of MBP and its precursor form, pre-MBP, via guanidinium chloride (GuHCl) titrations. Both proteins exhibited gradual unfolding: MBP began to unfold at ∼2.0 M GuHCl and was fully denatured by 4 M, whereas pre-MBP showed an earlier unfolding onset at 1 M but likewise reached full denaturation at 4 M. These results established key boundary conditions, providing critical reference points for interpreting intermediate states observed under non-equilibrium conditions (Fig. 1b and Supplementary note 1).

#### Real-time analysis of folding trajectories using NEXT-FRET

To analyze the stochastic, time-dependent smFRET data collected during MBP folding, we employed tvGMM^28^. This framework extends the conventional GMM by allowing the probability weights of each state to evolve over time, while keeping the Gaussian parameters—means and variances—fixed. This approach captures the time-dependent redistribution of molecular populations during folding, where state probabilities change over time while their structural characteristics remain constant. Temporal evolution of state probabilities was modeled using deterministic basis functions such as polynomials or B-splines, followed by a softmax transformation to ensure non-negative, normalized probabilities. This formulation enhances numerical stability and interpretability, allowing us to track the temporal evolution of states throughout the folding process. The tvGMM’s parameters were estimated via an adapted Expectation-Maximization (EM) algorithm^28,29^ which iteratively estimates both the time-dependent mixing weights and the parameters of the Gaussian components. The NEXT-FRET framework enables precise identification of transient folding intermediates and provides high-resolution insight into dynamic molecular transitions at the single-molecule level (Fig. 2a and Supplementary note 2).

**Fig. 2.**
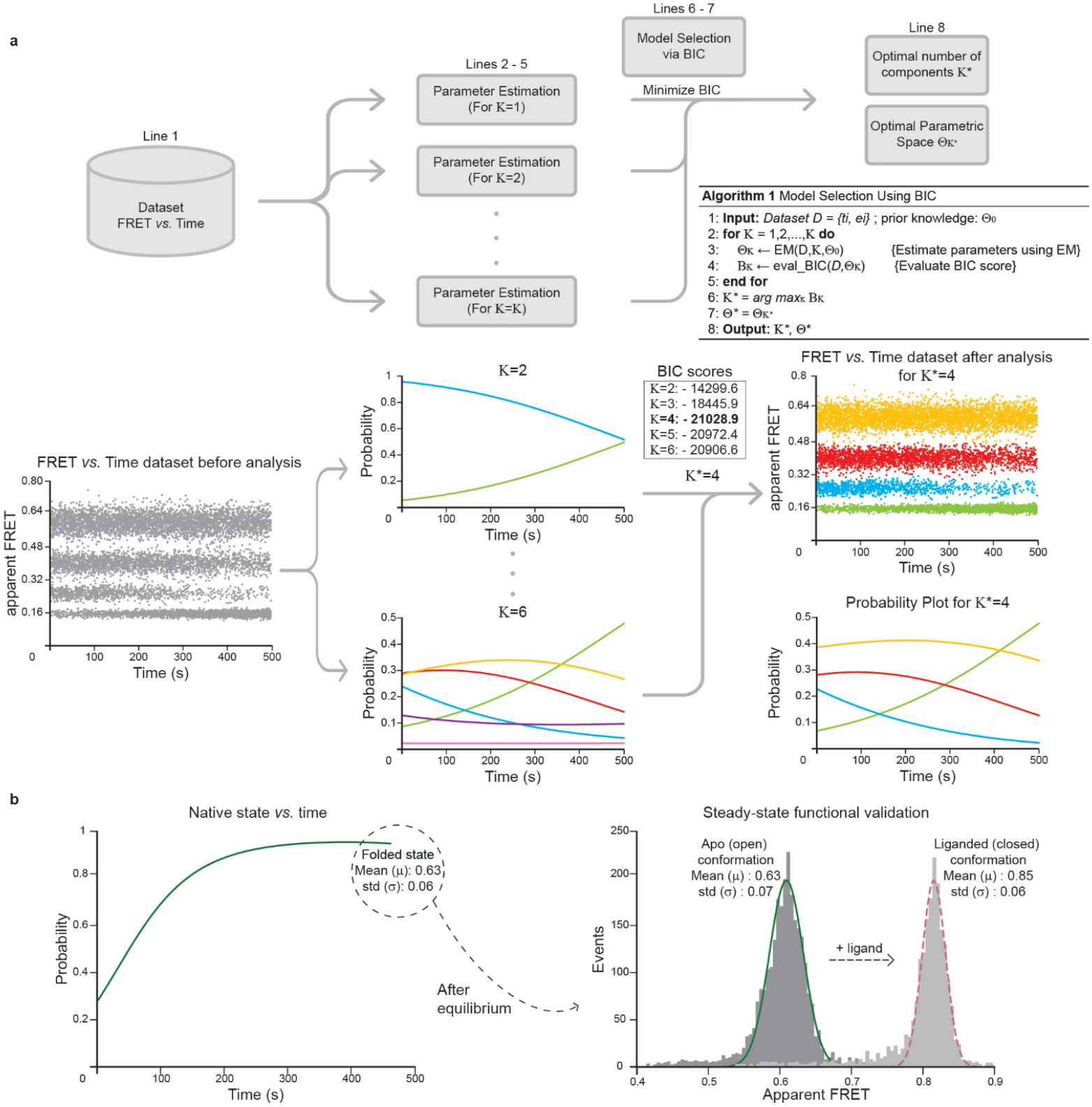
Time-dependent analysis of smFRET trajectories using NEXT-FRET. **a**, *Top panel*: The core algorithm is illustrated. Apparent FRET efficiency *vs*. time data analyzed to identify folding intermediates and their probabilities. *Bottom panel*: Application of NEXT-FRET to synthetic data. Parameters are estimated across models, with the optimal state number selected by BIC. The probability plot shows state likelihoods over time. **b**, NEXT-FRET analysis of experimental non-equilibrium smFRET data for MBP refolding. The apo-open state becomes increasingly populated after dilution from 6 M GuHCl. Final FRET efficiencies align with equilibrium values. Functional integrity is confirmed by maltose-induced transition to the closed conformation.

In addition to modeling time-varying probabilities, the framework incorporates two critical features: the integration of prior knowledge and model selection based on the Bayesian Information Criterion (BIC)^30^. Prior information—on both the temporal evolution of state probabilities and the parameters of the Gaussian components introduces regularization into the inference process. This Bayesian approach mitigates potential issues related to overfitting or insufficient sample sizes by leveraging prior knowledge from expert domain insights, thereby enhancing the reliability and robustness of parameter estimates. Moreover, utilizing BIC, the model balances complexity against goodness-of-fit, systematically comparing different number of possible intermediates and selecting the model that best minimizes the information criterion (Supplementary note 2 and Supplementary table 1).

To initiate non-equilibrium folding, MBP was rapidly diluted (>10^6^-fold) from 6 M GuHCl into aqueous buffer. This eliminated residual denaturant and allowed refolding to proceed under near-physiological conditions. smFRET measurements commenced immediately after dilution, capturing real-time folding dynamics (Supplementary note 1). The analysis revealed a progressive enrichment of the native open state (Fig. 2b). Initially, the population was dominated by unfolded conformations, which gradually transitioned toward the native open state, ultimately reaching a plateau.

Importantly, the post-folding FRET efficiency matched that of the equilibrium apo-open state, confirming full recovery of the functional state (Supplementary table 2). Functional validation was further achieved by adding maltose to the refolded protein, which induced the expected transition to the closed conformation. This confirmed that the refolded proteins retained structural integrity and biological activity, demonstrating the reliability of our experimental setup (Fig. 2b).

### Benchmarking NEXT-FRET against existing folding techniques

Single-molecule approaches —including detection methods like smFRET and manipulation techniques such as optical tweezers^31,32^— have long been employed to investigate protein folding. However, no single-molecule study to date has successfully detected folding intermediates in wild-type MBP. To overcome this challenge, earlier investigation introduced a slow-folding MBP double mutant in the presence of GuHCl, detecting an intermediate in the mutant but not in the wild-type protein^33^. Similarly, force-based approaches have supported a two-state folding model for MBP, but lacked the temporal and spatial resolution to distinguish intermediate states or infer structural insights^34,35^. Among ensemble techniques, hydrogen-deuterium exchange mass spectrometry (HDX-MS) has detected a folding intermediate in wild-type MBP under near-native conditions (0.2 M GuHCl), following denaturation from 2 M GuHCl^36^. While this study confirmed the presence of an intermediate with native-like contact patterns, it could not attribute its structural identity.

In contrast, NEXT-FRET, enables real-time tracking of the complete folding landscape. Crucially, the integration of prior knowledge—such as FRET efficiencies of the native state from equilibrium controls (Fig. 2b)— reduces the number of single-molecule trajectories required per condition, enabling efficient experimental designs with only a few thousand molecules (Supplementary note 2).

Using NEXT-FRET, we resolved a previously undetected folding intermediate in wild-type MBP. This state exhibits a higher FRET efficiency than the native open conformation and resembles the closed conformation observed upon maltose binding (Supplementary table 2). It represents a distinct, transient intermediate along the folding pathway. By identifying and characterizing this state, our method surpasses the resolution of both HDX-MS and prior single-molecule studies, bridging the gap between ensemble-averaged and single-molecule observations (Fig. 3a).

**Fig. 3.**
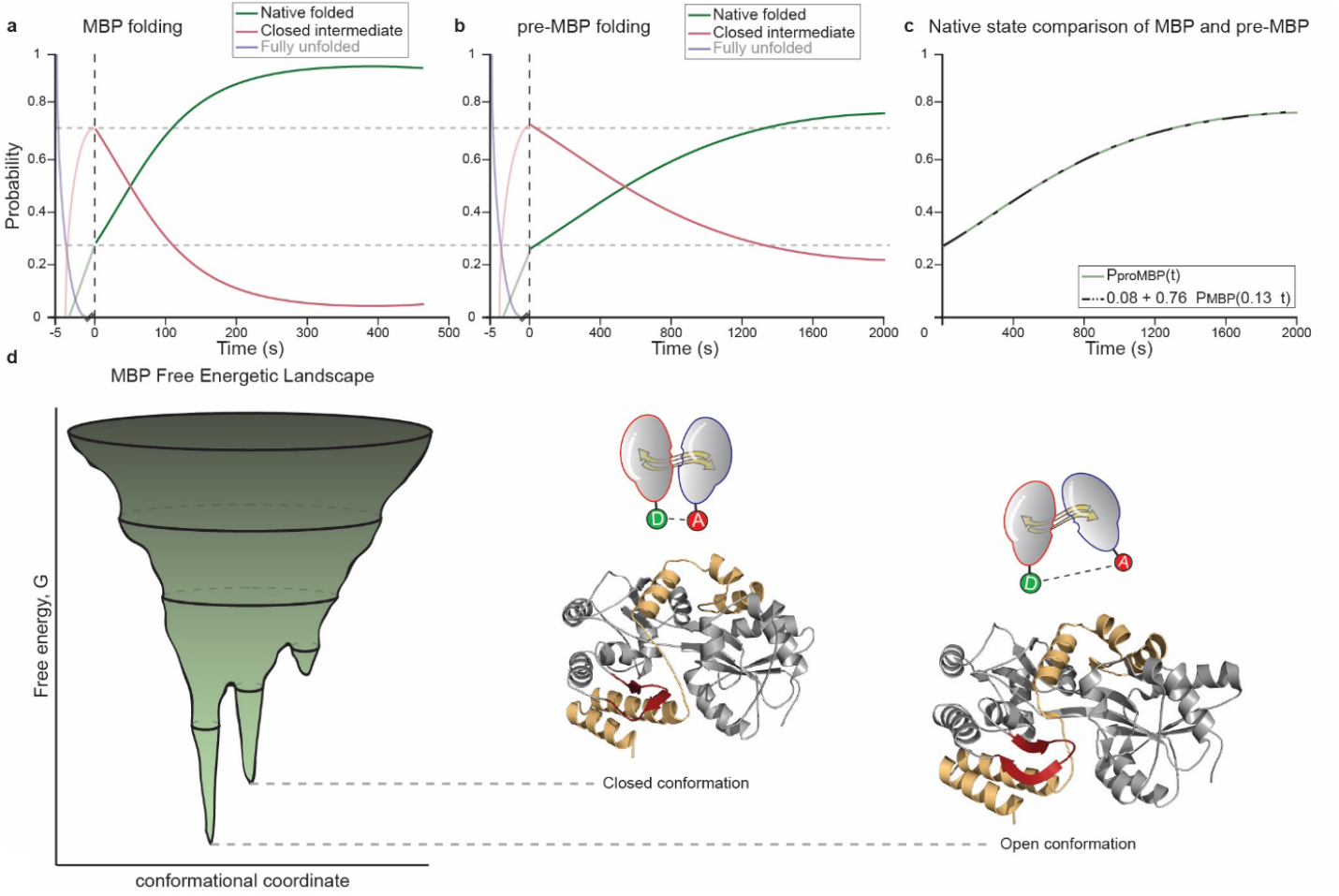
Characterization of the MBP folding intermediate and the effect of the signal peptide. **a–b**, Both MBP and pre-MBP traverse a shared folding intermediate before reaching the native open state. Faded traces indicate early transitions occurring faster than the measurement resolution. At time zero, ∼30% & 70% of molecules occupy the native and intermediate state respectively for both MBP and pre-MBP indicating that: **c**, SP delays only the intermediate-to-native transition by ∼7.5-fold and reduces folding efficiency to ∼80%. **d**, The intermediate resembles the closed, ligand-bound state. Key structural elements for conformational switching—C-terminal tail (yellow) and a dynamic regulatory region (red)—remain partially unfolded until transition to the open conformation^36^.

We next examined how the signal peptide (SP) modulated folding by analyzing pre-MBP. Remarkably, the same folding intermediate observed in wild-type MBP was also detected in pre-MBP (Fig. 3a,b). Folding in both proteins initiates from similar starting populations: ∼30% in the native open state and ∼70% in the intermediate, indicating that the SP does not delay the transition from the unfolded to the intermediate state. Instead, it selectively impedes the progression from the intermediate to the native open state. This reveals a precise modulatory role of the SP in shaping folding kinetics. Consistent with this view, folding of pre-MBP was delayed approximately 7.5-fold compared to MBP, although the overall pathway remained unchanged (Fig. 3c). This finding aligns with bulk studies showing that the SP interacts with the hydrophobic core of non-native proteins to delay folding^37^. Crucially, our single-molecule framework provides a direct visualization of this effect, offering unprecedented insight into how SPs reshape folding landscapes at the molecular level.

### Validating NEXT-FRET findings against the free-energy landscape of MBP

Protein folding proceeds along a rugged free-energy landscape (FEL), where biomolecules transition from high-energy unfolded states through intermediate states—key kinetic checkpoints—before reaching the low-energy, native state^25,38^.

Our smFRET data provide direct evidence that the closed conformation of MBP, typically observed upon ligand binding, functions as a bona fide folding intermediate. This finding offers structural and thermodynamic clarity consistent with FEL models. Prior studies have shown that the closed state occupies a higher free-energy level than the native open state^27^, aligning with our identification of this conformation as a stable, non-terminal intermediate (Fig. 3d).

Using HDX-MS, we previously demonstrated that ligand binding destabilizes the C-terminal tail (C-tail) of MBP, facilitating its transition to the closed conformation^27^. More recently, unpublished data from our group identified a second structural region (Fig. 3d, red) that cooperates with the C-tail to stabilize the open conformation (Muthahari Y.A, Gouridis G. et al., manuscript in preparation). Together, these two elements act as conformational restraints; both must be destabilized to enable acquisition of the closed state. While these studies elucidated the structural determinants of MBP’s conformational switching, they did not establish whether the closed state serves as an intermediate during folding.

Our present findings extend these observations. Walters et al.^36^ reported that the final regions to fold in MBP correspond to the same structural elements we previously identified: the C-tail (Fig. 3d, yellow) and the dynamic structural region (Fig. 3d, red). These regions form stabilizing contacts that maintain the native open state, as confirmed by recent work (Muthahari Y.A, Gouridis G. et al., manuscript in preparation). Moreover, Walters et al. found that the folding intermediate retains native-like contact patterns, supporting its structural resemblance to the closed conformation. This is fully consistent with our smFRET data, which identify the closed state as the intermediate bridging the unfolded and native open states. By integrating structural and thermodynamic evidence from both our current and prior studies, we now provide direct and conclusive evidence that the closed state is not merely a ligand-bound endpoint but a critical on-pathway intermediate in MBP folding.

In summary, our framework resolves transient folding intermediates with exceptional precision and identifies the closed conformation as a structurally and energetically significant intermediate in the FEL of MBP. By synthesizing thermodynamic and structural insights, we offer a unified model of MBP folding. This cohesive framework underscores the power of NEXT-FRET and its broad utility for dissecting the mechanisms of protein folding at high resolution.

### Performance evaluation: influence of chaperones and signal peptide on MBP folding

Understanding how molecular chaperones shape protein folding within the FEL is critical, as they modulate folding intermediates, prevent aggregation, and aid progression toward the native state^25^. NEXT-FRET enables high-precision detection of folding intermediates and provides a powerful platform to dissect how both chaperones and/or SPs influence the folding landscape. These insights extend beyond previous bulk and single-molecule studies, offering a detailed view of chaperone-assisted folding across dynamic, non-equilibrium conditions.

#### SecB: Stabilization of unfolded states

Bulk studies have shown that SecB reduces aggregation^39^, promotes translocation^40^, and delays folding by binding unfolded polypeptides^41,42^. Optical tweezers experiments previously demonstrated that SecB binds to unfolded MBP and stabilizes it in an extended state, without unfolding the native one^34^. Yet, these measurements lacked the resolution to identify distinct folding intermediates. Our analysis significantly extends these findings. We observe that SecB stabilizes two distinct unfolded states (Fig. 4a): a fully unfolded state characteristic of 6 M GuHCl (Fig. 1b), and a molten-globule-like state, matching the behavior at 2 M GuHCl (Fig. 1b)—consistent with previous reports on molten globules^43^. These two states are stabilized at the expense of the closed intermediate, which we have identified to be essential for native-state acquisition. Consequently, SecB delays MBP folding by selectively stabilizing early-stage intermediates (Supplementary table 2).

**Fig. 4.**
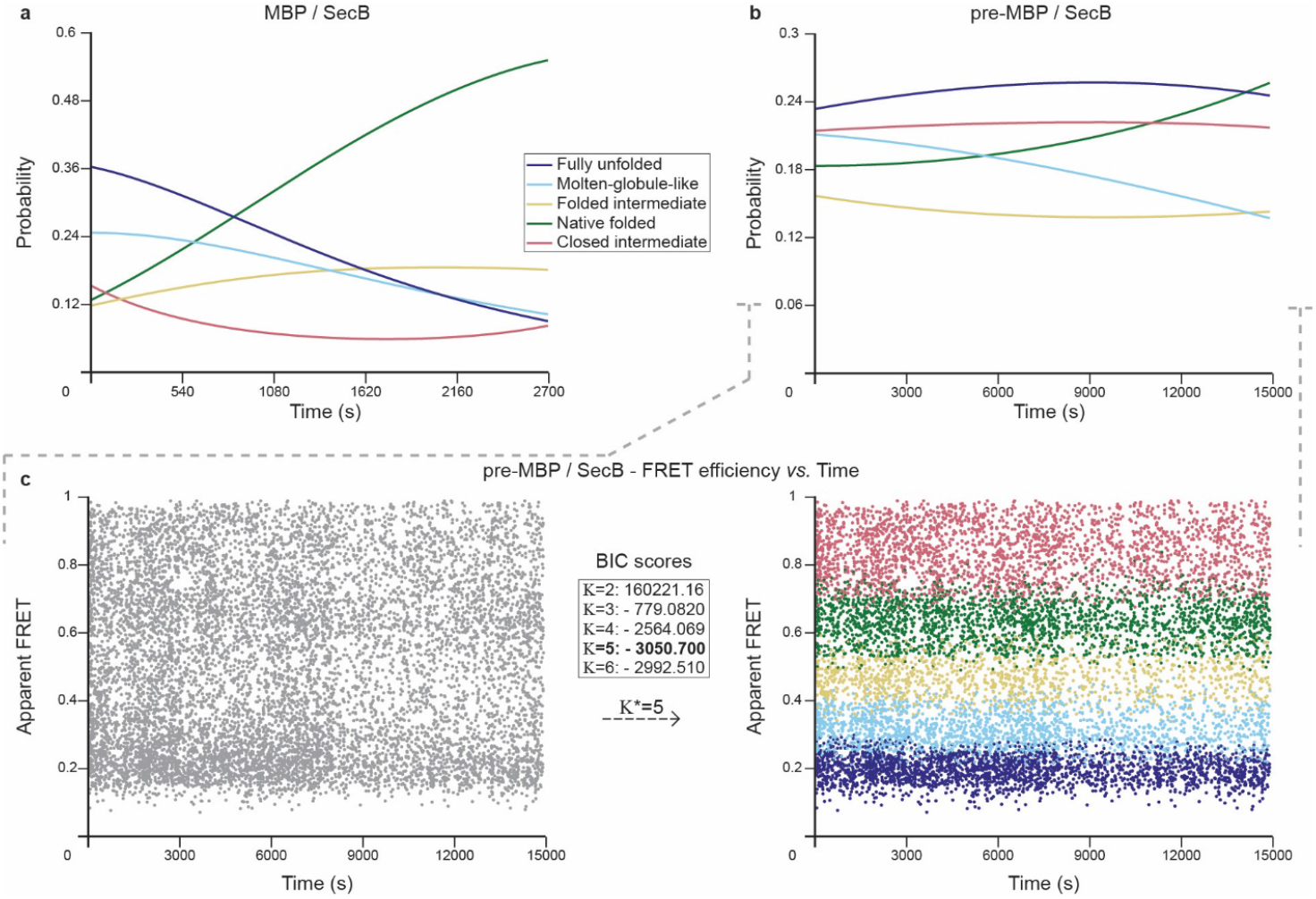
SecB selectively stabilizes early folding intermediates in MBP and pre-MBP. **a**, MBP folds through four states; SecB preferentially stabilizes two unfolded intermediates with low FRET efficiencies (dark/light blue). **b**, Pre-MBP follows a similar pathway, but the SP, together with SecB, enhances stabilization of the fully unfolded intermediate and delays folding progression. **c**, Time-resolved FRET traces from pre-MBP folding reveal five folding states, resolved using BIC-based selection within NEXT-FRET framework. This analysis captures the altered folding trajectory and highlights the synergy between SP and SecB in modulating folding dynamics at single-molecule resolution.

In pre-MBP, SecB exhibits distinct interaction dynamics (Fig. 4b). Although similar intermediates are detected, the majority of the population remains trapped in the fully unfolded state. This may reflect enhanced SecB affinity for pre-MBP or the presence of a signal peptide-induced energetic barrier that impedes folding progression. These results reveal a unique synergy between chaperone action and signal peptide modulation, shifting the folding equilibrium toward non-productive intermediates and highlighting their physiological relevance.

These conclusions are grounded on our advanced statistical analysis framework, which precisely resolves SecB-stabilized energetic wells. Traditional Gaussian fitting lacks the temporal sensitivity and adaptability needed to distinguish transient intermediates and fails to capture folding dynamins under non-equilibrium conditions (Supplementary figure 1). In contrast, NEXT-FRET enhances the resolution of smFRET data by allowing dynamic state probabilities while maintaining fixed means and variances for each folding intermediate. Integration of prior knowledge and model selection via BIC ensures a parsimonious yet expressive model, enabling robust inference of intermediate states without overfitting. As a result, our approach accurately identifies SecB-stabilized energetic wells, even within highly complex, non-equilibrium datasets (Fig. 4c).

#### Trigger Factor (TF): Stabilizing on-pathway intermediates

Unlike SecB, which primarily stabilizes unfolded states, TF preferentially stabilizes partially folded, on-pathway intermediates (Fig. 5a). Consistent with previous observations^35^, we found that TF promotes accumulation of a mid-FRET intermediate in MBP, while the fully unfolded state is absent. This intermediate—stabilized at the expense of the native state—represents a distinct, partially folded state that we refer to hereafter as the folded-intermediate (Supplementary table 2).

**Fig. 5.**
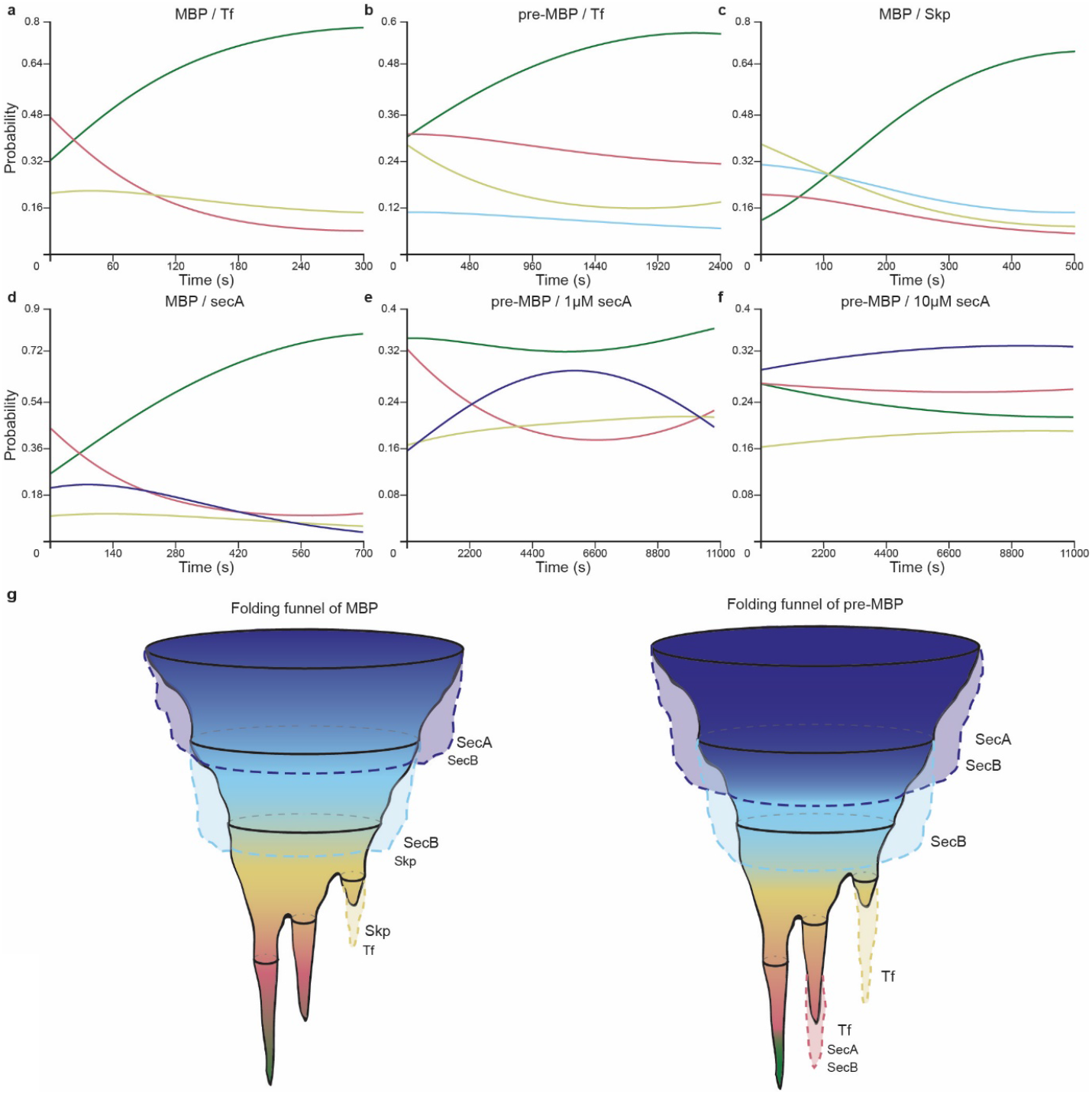
Chaperone-specific modulation on MBP and pre-MBP folding trajectories. **a–b**, TF traps folded-intermediate states, reducing native-state acquisition. **c**, Skp stabilizes both molten-globule-like and folded intermediates in MBP, modestly delaying folding. **d–f**, SecA progressively stabilizes non-native intermediates; in pre-MBP, this shift favors unfolded populations and prevents native folding at high concentrations. At sub-saturating (1µM) SecA concentrations, unfolded state increases at the expense of the closed one, consistent with SecA’s unfoldase activity. **g**, FEL illustrates chaperone-induced deepening of specific energetic wells. Stabilized intermediates are colored by FRET state. The SP broadens chaperone effects across the folding landscape, reinforcing kinetic barriers and reducing folding efficiency.

TF binding inhibits folding completion by kinetically trapping molecules in this intermediate, thereby reducing the fraction of molecules reaching the native state by ∼20%. This “freezing” effect illustrates TF’s ability to selectively modulate folding dynamics by stabilizing specific FEL wells along the folding pathway.

When the same analysis was extended to pre-MBP, both the folded-intermediate and a residual unfolded population were detected (Supplementary table 2). In this context, the signal peptide introduces an additional layer of modulation: the closed state is also prominently stabilized, alongside the folded-intermediate; collectively impeding efficient progression to the native open state. As a result, only ∼45% of molecules reach the native conformation in the combined presence of TF and the SP (Fig. 5b). These findings underscore TF’s mechanistic specificity—stabilizing on-pathway intermediates rather than unfolded states—and highlight how the SP reshapes folding trajectories by reinforcing kinetic traps that interfere with native-state acquisition.

#### Skp: Modulating folding through stabilization of diverse intermediates

Skp, a periplasmic chaperone traditionally associated with outer membrane proteins, has been shown to interact with MBP, stabilizing unfolded and partially folded states through hydrophobic and electrostatic interactions^44,45^. Our analysis indicates that Skp preferentially stabilizes the folded-intermediate state and, to a lesser extent, the molten-globule-like state, while reducing the population of the closed intermediate (Fig. 5c and Supplementary table 2). This shift in the folding landscape delays progression toward the native state, reducing the overall folding rate of MBP.

In the case of pre-MBP, Skp induced nonspecific aggregation, likely due to hydrophobic interactions between exposed residues and the experimental surface. As this behavior does not reflect physiologically relevant folding dynamics and compromises data reliability, we did not pursue further analysis on Skp’s effect on pre-MBP.

#### SecA: An active modulator of folding and unfolding via stabilization of non-native states

SecA, the cytoplasmic motor ATPase of the Sec translocon, is essential for post-translational protein translocation across the inner membrane^46^ and has been proposed to function as a cellular unfoldase^47^. Our smFRET data demonstrate that SecA profoundly alters MBP’s folding landscape by stabilizing both the fully unfolded state and the folded-intermediate state (Fig. 5d and Supplementary table 2). Over time, these states gradually depopulate as molecules transition to the native open conformation. However, this transition occurs at a markedly slower rate than in the absence of SecA, indicating that SecA imposes a kinetic delay by stabilizing non-native intermediates.

Under physiologically relevant conditions, pre-MBP exhibits a distinct folding response to SecA. Specifically, the percentage of the closed intermediate state decreases, while the fully unfolded state becomes increasingly dominant, suggesting that SecA actively unfolds intermediate states (Fig. 5e). Meanwhile, the folded-intermediate and native open states remain relatively stable, revealing a differential modulation of the folding pathway in pre-MBP compared to MBP.

At saturating SecA concentrations (10 µM)^48^, we observe broad stabilization of all folding intermediates—including the folded-intermediate and closed states—while the native open state is strongly suppressed (Fig. 5f). Although SecA is known not to bind fully folded proteins^48^, its interaction with intermediate states is sufficient to shift the folding equilibrium toward non-native states. Over time, the fully unfolded state becomes increasingly populated, consistent with an ATP-independent unfoldase function.

These findings support a model in which SecA selectively engages and stabilizes non-native intermediates to regulate folding dynamics. By reshaping the FEL and promoting accumulation of non-native states, SecA effectively biases the folding-unfolding equilibrium away from the native state. This dual function—as both a translocation motor and a folding regulator—underscores SecA’s central role in protein quality control and export.

## Discussion

Most *in vitro* folding studies have focused on small, single-domain proteins, simplifying analysis but overlooking the complexity of multidomain systems. Approximately 28% of proteins contain discontinuous domains^49^, which inherently reduce folding efficiency but enhance regulatory and functional versatility^50,51^. To benchmark analyses of folding mechanisms under physiologically relevant conditions, we selected MBP, a well-studied two-domain substrate-binding protein (SBP) with a discontinuous fold. SBPs serve as extracellular solute receptors partnering with ATP-Binding Cassette (ABC) transporters^52^ and are essential for nutrient uptake, cell-cell communication, and virulence in diverse pathogens^53–59^.

Beyond elucidating folding landscapes, our findings offer a novel conceptual framework for therapeutic intervention. We demonstrate that chaperones, such as SecB, can selectively stabilize folding intermediates, effectively arresting folding progression. The emerging paradigm of “therapeutic folding arrest”^60^ bypasses traditional occupancy-based inhibition, instead trapping proteins in non-functional states prior to native-state formation. Additionally, the mechanisms we identified (Fig. 3-5) highlight the central role of chaperones in maintaining proteostasis by controlling the conformational states available to proteins, preventing aggregation, and ensuring cellular health^25^. Disruptions in chaperone-mediated folding landscapes have been increasingly linked to the molecular pathogenesis of neurodegenerative and other conformational diseases, underscoring the biomedical significance of resolving these intermediate states^25^. Conformational entrapment strategies open new avenues for targeting highly dynamic proteins, those lacking well-defined binding pockets, or whose native clefts are occupied by high-affinity natural ligands—hallmarks of traditionally “undruggable” targets^61–65^. Moreover, such strategies may help preclude off-pathway aggregation events, such as amyloid formation, by intercepting metastable intermediates before misfolding occurs^25,66^.

SBPs exemplify such challenging targets. Despite their centrality to microbial fitness—including several WHO-priority pathogens—SBPs have been overlooked in drug discovery due to the absence of stable pockets in their native states resulting from the Venus flytrap–like ligand capture mechanism^67^. By identifying folding intermediates as kinetic vulnerabilities, our results open new avenues to exploit these transient states through rational design of small molecules or chaperone-mimicking biologics capable of folding arrest.

NEXT-FRET, by revealing such intermediates under non-perturbative, near-physiological conditions, offers a scalable platform for screening molecules that modulate protein folding dynamics. This capability significantly expands opportunities for therapeutic targeting of conformational plasticity in folding-defective precursors, aggregation-prone proteins implicated in proteostasis collapse, and conformationally regulated signaling proteins involved in disease pathways.

In conclusion, NEXT-FRET enables real-time, solution-phase tracking of protein folding trajectories, bridging the methodological gap between ensemble and single-molecule studies. Future integration with rapid-mixing technologies can extend temporal resolution to early folding events; e.g., within the first ∼5 s post-refolding initiation (Fig. 3a,b). The simplicity, robustness and adaptability of NEXT-FRET render it ideally suited for dissecting non-equilibrium biomolecular processes in *in vitro* reconstituted systems, thereby expanding the chemical biology toolkit towards non-traditional, dynamic therapeutic targets.

## Methods

### Derivation of the Time-varying Gaussian Mixture Model in NEXT-FRET

Consider an smFRET experiment beginning at time *T*_0_ and ending at final time *T*_1_, where 0 ≤ *T*_0_ *< T*_1_. During this interval, measurements are recorded as pairs (*e, t*), where *t* ∈ [*T*_0_, *T*_1_] denotes the detection time of a molecule, and *e* := *e*(*t*) corresponds to the associated apparent FRET efficiency value. The joint probability density of (*e, t*) factorizes as

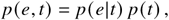

where *p* (*t*) is the probability density of detection times, and *p* (*e*|*t*) is the conditional probability of apparent FRET efficiencies given the detection time *t*. The distribution *p* (*t*) depends on the experimental conditions, including the number of molecules that pass through the confocal volume over time, and is not associated to the actual folding dynamics. In contrast, *p* (*e*|*t*) encodes the time-dependent behavior of apparent FRET efficiencies and is the main focus of our modeling effort, as it provides direct insight into the temporal evolution of the folding process.

The measured apparent FRET efficiency is directly determined by the distance between donor and acceptor fluorophores ^68^, which is uniquely specified by the molecule’s conformational state. This conformational state is modeled as a discrete latent random variable *Z*, taking values in the set {1, …, *K*}, where *K* denotes the total number of identifiable conformational states. The variable *Z* is latent, as it is not directly measured. Central to our analysis is the fact that, once *Z* is known, the apparent FRET efficiency is conditionally independent of the detection time. Due to the non-equilibrium nature of the protein folding process, the distribution of the latent variable *Z* varies over time. Specifically, the probability of a molecule occupying conformational state *k* at time *t* is denoted by *p*_*k*_ (*t*) := *p* (*Z* = *k* |*t*), for *k* = 1, …, *K*. These time-dependent probabilities model how the occupancy of different folding intermediates evolves throughout the experiment.

Assuming that each conformational state *Z* = *k* corresponds to a distinct donor-acceptor distance, we define the distribution of the apparent FRET efficiency conditioned on the *k*-th state as *p*_*k*_ (*e*) := *p* (*e*|*Z* = *k*). Then, using the law of total probability and incorporating the aforementioned conditional independence property, the conditional distribution of apparent FRET efficiency given the detection time *t* can be expressed as:

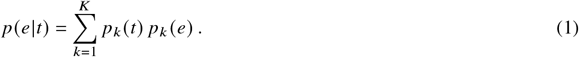

This probabilistic model represents a mixture distribution with time-varying mixing weights, *p*_*k*_ (*t*). It effectively captures the non-equilibrium dynamics of the folding process, while assuming that the emission distribution for each folding intermediate, *p*_*k*_ (*e*), remains constant over time.

We now proceed with the parametrization of the two conditional distributions. Given the low concentrations (in the *pM* range) employed in smFRET experiments, the likelihood of two molecules simultaneously entering the excitation volume is negligible. As a result, each single molecule is assumed to adopt one and only one of the *K* distinct folding intermediates. Additionally, although photon emission is inherently stochastic and governed by a Bernoulli process, the FRET efficiency is computed from the total number of detected photons. Since this count aggregates many independent emission events, the law of large numbers applies, yielding a distribution that is approximately Gaussian. Consequently, the distribution of apparent FRET efficiency conditioned on each conformational state, denoted by *p*_*k*_ (*e*), is modeled as a Gaussian distribution with mean value *μ*_*k*_ and variance 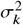. This leads to the formulation of a probabilistic latent model described as a *time-varying Gaussian Mixture Model (tvGMM)*, in which the mixing weights *p*_*k*_ (*t*) vary with time capturing the folding dynamics, while the parameters of the Gaussian components remain constant.

We adopt a semi-parametric approach to model the probability that a single molecule is in conformational state *k* at time *t, p*_*k*_ (*t*). This choice offers significantly greater flexibility compared to fully parametric models. Specifically, we define a softmax-based formulation for the *K* states, where the probability of the *k*-th state is given by:

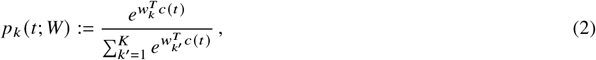

where 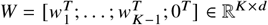 represents the parameter matrix while *c*(*t*) ∈ R^*d*^ denotes a vector of deterministic basis functions. This modeling approach allows us to represent each probability of a conformation state as a weighted linear combination of the basis functions. To ensure identifiability and numerical stability in parameter estimation, the last row of *W* is fixed to zero. This constraint resolves the inherent redundancy of the softmax function since the resulting probabilities must always sum to one. The vector function *c*(*t*) : R_+_ → R^*d*^ consists of deterministic basis elements designed to capture both constant and oscillatory behavior. In particular, *c*(*t*) may include low-order polynomial terms, Fourier components (e.g., sin(2 πω*t*), cos(2πω*t*) for various frequency values, ω), and B-spline functions, which together provide a flexible representation of non-linear temporal variation. The resulting model for *p*_*k*_ (*t*) can be also interpreted as a shallow artificial neural network with no hidden layers: the input *c*(*t*) is linearly transformed by *W*, and the output is passed through a softmax activation function to yield a valid probability distribution.

Overall, the parametrized version of Eq. (1) takes the form:

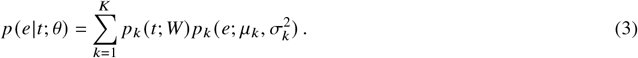

where 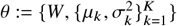 is the parameter set of tvGMM whose total count is |θ | = *K* (*d* + 1).

### Parameter Estimation and Expectation Maximization Algorithm

Each single molecule is identified by an index i where i ranges from 1 to N where N is the total number of measured molecules. The dataset D = {(*e*_1_, *t*_1_), …, (*e*_N_, *t*_N_)} consists of apparent FRET efficiency measurements *e*_i_, each recorded at a corresponding detection time *t*_i_ for the i-th molecule. Based on the standard modeling assumption that single molecule conformations are independent, the corresponding log-likelihood function is expressed as:

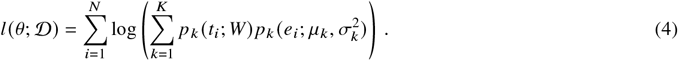

Maximum Likelihood Estimation (MLE) ^69–71^ is a foundational approach for estimating the parameters of statistical models by maximizing the above log-likelihood of observed data. It has been extensively applied in the analysis of single-molecule datasets^72,73^, including smFRET measurements ^74–76^. MLE operates without incorporating prior knowledge about the parameters being estimated. However, in proteins, certain conformational states –such as the native folded and the liganded ones– are well characterized, with their apparent FRET statistical properties experimentally determined. Incorporating this prior information can improve estimation, especially when model assumptions are not fully satisfied or when data availability is limited. To improve the robustness of the parameter estimation and subsequently the biological interpretability of the estimates, we incorporate prior knowledge into the estimation approach using a Bayesian framework for some of the parameters. Accordingly, we impose prior knowledge to the parameters of the Gaussian components using conjugate prior distributions ^77,78^. The conjugate prior probability for the mean value of the *k*-th component is Gaussian and it takes the form:

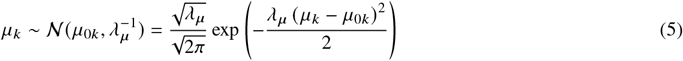

where *μ*_0*k*_ is the mean of the prior distribution for the *k*-th state while λ_*μ*_ is the inverse variance and it is interpreted as the strength of the prior (e.g., larger values for λ_*μ*_ implies stronger prior). As a conjugate prior for the variance parameter 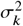 of the *k*-th Gaussian component, we employ the Inverse Gamma distribution, expressed as:

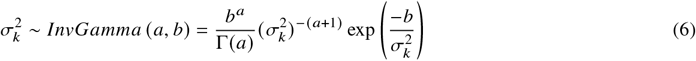

with 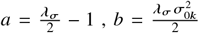 where 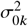 corresponds to the mode (i.e., the value that maximizes the probability density function) of the Inverse Gamma distribution while λ _σ_ is the respective strength of the prior.

The inclusion of conjugate priors modifies the standard log-likelihood to the following extended form:

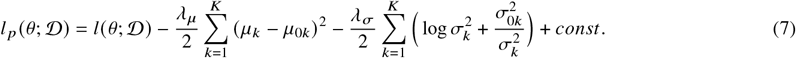

whose maximization leads to Maximum A Posteriori (MAP) estimation of the parameters.

The optimization of Eq. (7) with respect to θ is non-linear and it requires an iterative approach. The standard estimation procedure for GMMs^29^ can be extended to the time-varying case with priors, leading to a novel variant of the Expectation-Maximization (EM) algorithm. The EM algorithm is a pivotal iterative method that is widely used for parameter estimation in probabilistic models, especially when data is incomplete or involves latent variables ^29,79,*80*^. *It iteratively alternates between two steps: the Expectation step (E-step), which computes the expected log-likelihood with respect to the current parameter estimates, and the Maximization step (M-step), which updates the parameters by maximizing this expected log-likelihood. This process is repeated until convergence*.

*In the E-step, we compute the posterior probability of the* i-th sample being generated by the *k*-th state, denoted by γ_i,*k*_ = γ_i,*k*_ (θ) := *p* (*Z* = *k* |*e*_i_, *t*_i_; θ), also known as *responsibility*. It is calculated using the Bayes’ theorem as:

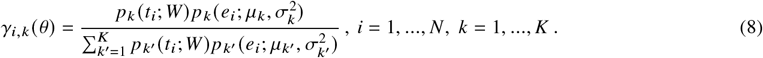

Based on the *responsibilities*, a lower bound of the log-likelihood *l* (θ; D) is constructed as the expectation of the complete-data log-likelihood, evaluated at a fixed parameter value 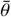:

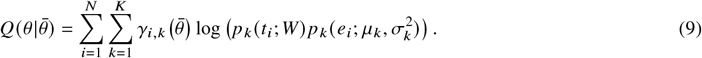

In the M-Step, the log-likelihood in Eq. (7) is replaced by 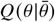 from Eq. (9), which is then maximized with respect to the current parameter estimate 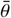 to obtain an updated estimate 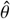:

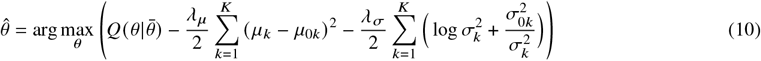

Since Eq. (10) can be optimized independently with respect to *W* and the Gaussian parameters, the procedure is divided into two sub-steps. The first involves maximizing with respect to *W* :

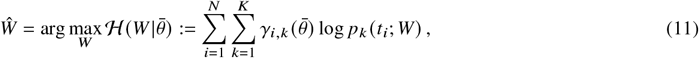

which results in an updated estimate for *W*. To perform this optimization, we employ the Newton-Conjugate Gradient (Newton-CG) method ^81^, a second-order optimization technique that leverages both gradient and Hessian information to efficiently navigate the parameter space. This method is particularly suited for smooth, potentially high-dimensional objective functions, enabling robust and rapid convergence.

The second sub-step involves maximizing Eq. (10) with respect to the parameters of the Gaussian components, *μ*_*k*_ and 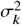, for each *k*, which leads to the following closed-form updates:

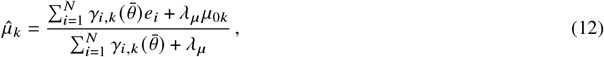

for the *k*-th mean while for the *k*-th variance it is given by:

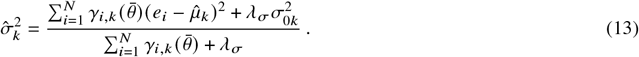

The E- and M-steps are iteratively repeated until convergence, which is typically assessed by monitoring the change in the expected log-likelihood and terminating when it falls below a predefined threshold ε. The complete procedure is outlined in Algorithm 1. The EM algorithm is initialized by setting the weight matrix *W* = 0, which yields uniform time-varying mixing weights at the first iteration. The means *μ*_*k*_ of the Gaussian components are initialized by partitioning the range of apparent FRET efficiency values into *K* equally spaced segments. The variances 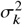 are initialized to a fixed constant across all components. Finally, it is important to note that while the EM algorithm is not guaranteed to convergence to a global maximum, nevertheless, it is guaranteed to produce a non-decreasing sequence of log-likelihood values across iterations.

#### Algorithm 1

Expectation Maximization for tvGMM parameter estimation

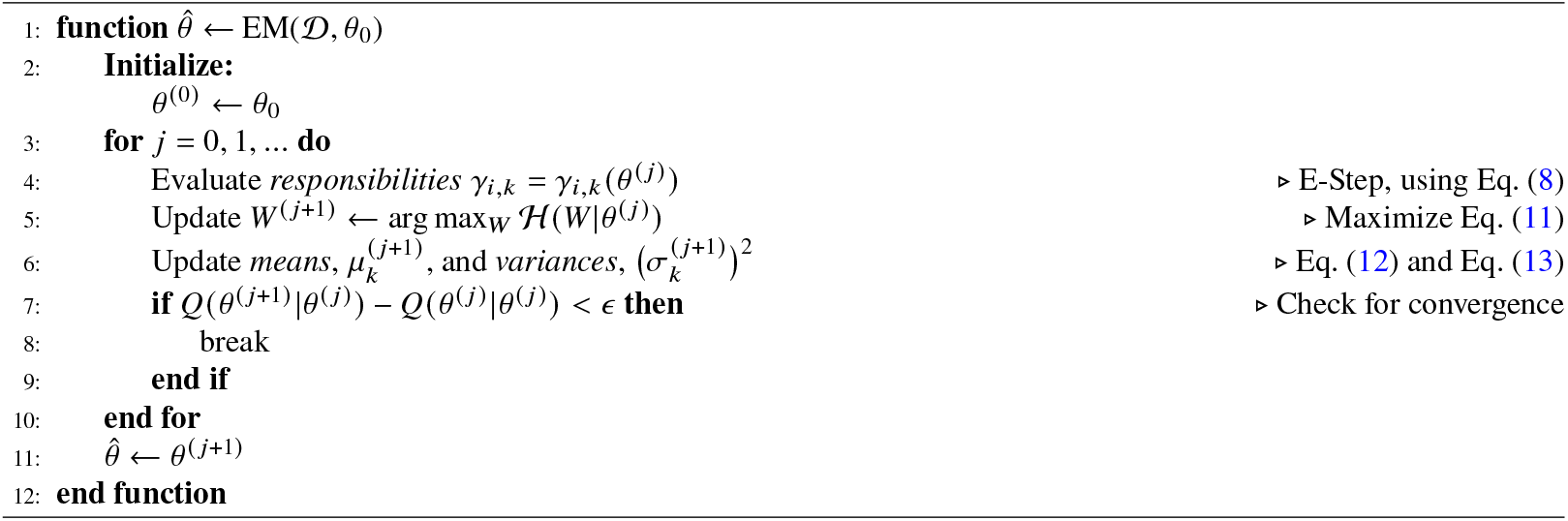

### Selecting the Number of Folding Intermediates

Within both the MLE and MAP estimation frameworks, model selection is essential for determining the appropriate number of latent conformational states. The Bayesian Information Criterion (BIC)^30, 82^ *offers a principled means for this by incorporating a penalty term that accounts for model complexity. By evaluating BIC across varying numbers of states K*, we select the model that best balances fit and simplicity. In the proposed tvGMM case, the BIC is defined as:

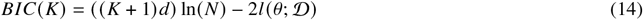

where *l* (θ; D) is the log-likelihood given by Eq. (4).

In our analysis, the number of basis functions *d* (i.e., the dimension of *c*(*t*)) is chosen to be relatively small, typically between 2 and 5. This low-dimensional representation implicitly enforces smoothness in the time-varying mixing weights, serving as a form of regularization. Additionally, the temporal resolution of the smFRET measurements and the gradual evolution of the folding dynamics limit the need for larger values of *d*. The effectiveness of the tvGMM parameter estimation and model selection strategies is quantitatively validated through synthetic data experiments, as detailed in the Supplementary Notes.

### Matching the kinetics of two folding experiments

To compare folding trajectories across different experimental conditions, we introduce a transformation to match the respective time-varying state probability functions. Our objective is to align *p*_*k*_ (*t*; *W*) from the first experiment with the corresponding distribution 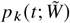 from the second. Dropping the *k* dependence from the notation, the transformation is defined as:

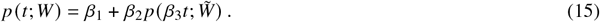

In Eq. (15), the parameter β_1_ serves as an intercept term that captures differences in the initial percentage of a given folding intermediate at time *T*_0_; values near zero indicate similar initial state percentages between experimental conditions. The parameter β_2_ scales the overall magnitude of the probability curve. When β_1_ ≈ 0, β_2_ approximates the ratio of plateau level state probabilities for a specific intermediate, with a value close to one implying comparable folding efficiencies (number of molecules reaching the native state). Lastly, β_3_ acts as a temporal scaling factor. Under the assumption that β_1_ ≈ 0, deviations of β_3_ from unity reflect differences in the folding rates between the two experimental conditions—values less than one indicate temporal dilation (slower folding), while values greater than one correspond to temporal contraction (faster folding). Overall, this transformation allows for a direct and interpretable comparison of the folding time-dependent trajectories between the two experiments for a given folding intermediate.

Finally, the coefficients β_1_, β_2_, β_3_ are inferred by solving the following non-linear least-squares problem:

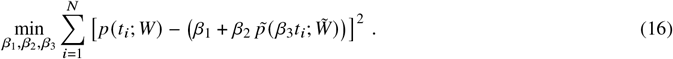

Due to the non-linearity of the temporal scaling parameter β_3_, we utilize Newton’s method which is a first-order iterative optimization algorithm for the estimation of the coefficients.

## Data availability

All data supporting the findings of this study are available within the main text and Supplementary Information. Data acquisition and analysis were performed using established, peer-reviewed protocols and publicly available software tools or servers, as referenced throughout the manuscript.

## Code Availability

The code used for NEXT-FRET is available at https://github.com/andrewkof/NEXT-FRET. All other analyses were conducted using standard, publicly available software tools, as cited in the manuscript. The complete codebase will be made publicly available upon acceptance.

## Acknowledgments

We thank T. Cordes for providing software for data acquisition, analysis of ALEX data, and expert guidance in assembling the smFRET confocal setup. This work was supported by the Emblematic Action Brain Precision Grant TAEDR-0535850 (G.G.); a Start-up Grant from the Institute of Molecular Biology and Biotechnology (G.G.); the Theodore Papazoglou FORTH Synergy Grant 2022 (G.G. & Y.P.); the H2020 FETOPEN grant Dynamic (GA-863203); the H2020 FET Innovation Launchpad grant Wavelens (GA-101035014); and the HFRI Fellowship number 488.

## Author contributions

Project Initiation: YP, GG, MdB Conceptualization: CS, AK, MjB, YP, GG; Investigation: CS, AK,MdB, GZ, GG; Formal Analysis: CS, AK, YP, GG; Visualization: CS, AK, YP, GG; Infrastructure assembly: GZ, MM, KM Supervision: GZ, YP, GG; Writing-original draft: CS, AK, YP, GG; Writing-review & editing: CS, AK, YP, GG

## Corresponding contributions

Correspondence and requests of materials should be addressed to Yannis Pantazis and Giorgos Gouridis.

