## Supplementary Information for "NEXT-FRET: A solution-based smFRET platform to resolve folding intermediates under native conditions"

### Supplementary Note 1

##### *Reagents*

DNA-modifying enzymes used in molecular biology were sourced from NEB. Unless specified otherwise, all chemicals were obtained from Sigma Aldrich. Alexa Fluor 555 and Alexa 647 were supplied by ThermoFisher. (±)−6-Hydroxy-2,5,7,8-tetramethylchromane-2-carboxylic acid (Trolox) came from Merck. All chromatography materials were provided by GE Healthcare, and Ni-NTA resin was from Qiagen.

##### *Gene isolation and protein expression*

secA gene (UniProt: P10408) and secB gene (UniProt: A8A674) were isolated from the genome of *Escherichia coli* K12. Primers introduced *Nde*I and *Hind*III restrictions sites, and the gene product was sub-cloned into the pET16b vector (Novagen).

skp gene (UniProt: P0AEU7) and tig gene (UniProt: P0A850) were isolated from the genome of *Escherichia coli* K12. Primers introduced *Nde*I and *Xho*I restrictions sites, and the gene product was sub-cloned into the pET16b vector (Novagen).

malE gene (UniProt: P0AEX9) was isolated from the genome of *Escherichia coli* K12 with and without the signal peptide (MBP and pre-MBP). Primers introduced *Nde*I and *Hind*III restrictions sites, and the gene products were sub-cloned into the pET20b vector (Merck).

BL21 DE3 cells (F– ompT gal dcm lon hsdSB(rB–mB–) λ(DE3 [lacI lacUV5-T7p07 ind1 sam7 nin5]) [malB+]K-12(λS)) was used to over-express MBP-His$_6$, His$_6$-secA, secB-His$_6$, Skp-His$_6$ and Trigger factor-His$_6$. Cells harboring plasmids expressing the protein were grown in LB medium (37ºC; $OD_{600\ nm}$=0.8) and protein overexpression was induced by IPTG (0.5 mM; for 4 hours). In the case of pre-MBP, prior to induction, 4mM sodium azide were added to the media to block the secYEG system from exporting pre-MBP to the periplasm. Pre-MBP was induced with IPTG (1mM; for 3 hours).

##### *Purification of MBP*

Harvested cells of MBP were diluted in 50 mM Tris-HCl, pH = 8; 1 M KCl, 10% glycerol; 10 mM Imidazole; 1 mM Dithiothreitol (DTT); 2 mM PMSF and subsequently lysed by French press (800 psi; 3-4 rounds). After centrifugation at 50,000 g for 30 min (4 ºC; Sorval), the soluble material was loaded on a Ni-NTA resin (Qiagen). Bound proteins were washed (50 mM Tris-HCl, pH = 8; 1 M KCl, 10% glycerol; 10 mM Imidazole; 1 mM DTT; and 50 mM Tris-HCl, pH=8; 50 mM KCl, 10% glycerol; 30 mM Imidazole; 1 mM DTT sequentially) and then eluted (50 mM Tris-HCl, pH=8; 50 mM KCl, 10% glycerol; 300 mM Imidazole; 1 mM DTT). Protein fractions were pooled (supplemented with 5 mM EDTA; 50 mM DTT), concentrated (Vivacell-Sartorious; Amicon-Millipore), dialyzed (50 mM Tris-HCl, pH=8; 50 mM KCl, 50% glycerol; 10 mM DTT), aliquoted and stored at -80ºC.

***Purification of pre-MBP***

Cell pellets of pre-MBP were resuspended in 50 mM Tris-HCl, pH = 8; 500 mM NaCl; 5 mM $MgCl_2$; 5% glycerol; PMSF (1 mM) and DTT (10 mM). Cells were lysed by sonication. The lysate was centrifuged (18,000 rpm, 30 min, 15 ºC) to separate membrane and soluble fractions. The membrane fraction was resuspended in 50 mM Tris-HCl, pH = 8; 500 mM NaCl; 5 mM $MgCl_2$; 8 M Urea; 5% glycerol using a 2mL Dounce homogenizer. The sample was then diluted to 60 mL with the same buffer and stirred at 4 ºC. Membrane-solubilized material was clarified by centrifugation at 50,000 g for 30 min (4 ºC; Sorval), and the supernatant was diluted to a final buffer concentration of 50 mM Tris-HCl, pH = 8; 500 mM NaCl; 5 mM $MgCl_2$; 6 M Urea; 5% glycerol before being loaded onto a pre-equilibrated Ni-NTA resin (Qiagen). Bound proteins were washed sequentially with buffer of 50 mM Tris-HCl, pH = 8; 500 mM NaCl; 5 mM $MgCl_2$; 6 M Urea; 5% glycerol and with buffer of 50 mM Tris-HCl, pH = 8; 50 mM NaCl; 5 mM $MgCl_2$; 6 M Urea; 5% glycerol). UREA concentration was reduced by washing the column with the same buffer reducing the concentration to 4M, 2M, 1M and finally got rid of UREA entirely.  Elution was performed using 50 mM Tris-HCl, pH = 8; 50 mM NaCl; 100 mM Imidazole; 5 mM $MgCl_2$; 5% glycerol. Eluted protein fractions were dialyzed (50 mM Tris-HCl, pH=8; 50 mM NaCl, 50% glycerol; 10 mM DTT), aliquoted and stored at -80ºC.

***Chaperone purification***

Harvested cells of Chaperones, including secA, secB, Trigger factor (TF) and Skp were diluted in 50 mM Tris-HCl, pH = 8; 500 mM NaCl, 5% glycerol and 5 mM Imidazole and subsequently lysed by sonication (describe setup). After centrifugation at 50,000 g for 30 min (4 ºC; Sorval), the soluble material was loaded on a Ni-NTA resin (Qiagen). Bound proteins were washed (50 mM Tris-HCl, pH =

8; 500 mM NaCl, 5% glycerol; 5 mM Imidazole and 50 mM Tris-HCl, pH=8; 50 mM NaCl, 5% glycerol; 30 mM Imidazole; 1 mM DTT sequentially) and then eluted (50 mM Tris-HCl, pH=8; 50 mM NaCl, 5% glycerol; 300 mM Imidazole). Protein fractions were pooled and dialyzed (50 mM Tris-HCl, pH=8; 50 mM NaCl, 50% glycerol), aliquoted and stored at -80ºC. Proteins were further purified by size-exclusion chromatography (Superdex 200, GE Healthcare).

***Protein labelling***

MBP labeling was performed as previously described[27]. Following structural analysis of the open and closed x-ray crystal structures of MBP (PDB IDs: 1OMP and 1ANF, respectively), residues 36 and 352 were mutated to cysteines. Both MBP-$His_6$ and pre-MBP-$His_6$ (retaining its signal peptide) were immobilized on Ni-NTA resin (Qiagen) in the presence of 1 mM DTT to maintain the cysteines in their reduced state. The resin was incubated at 4°C for 2–8 hours in 50 mM Tris-HCl (pH 7.4) supplemented with 50 nmol of Alexa Fluor 555 and Alexa 647 (ThermoFisher). Unbound fluorophores were removed through extensive washing, and the labeled protein was further purified by size-exclusion chromatography (Superdex 200, GE Healthcare) to enrich the double-labeled fraction and eliminate potential aggregates. For folding experiments, MBP and pre-MBP were labeled in the presence of 6 M GuHCl in all buffers throughout the labeling, washing, and elution steps to ensure complete protein unfolding. In this case, size-exclusion chromatography was not applicable. Labeling efficiency for all proteins exceeded 80%.

***Solution-based smFRET and ALEX***

ALEX experiments were carried out at 25-50 pM of double-labelled protein in the appropriate buffer (50 mM Tris-HCl, pH 7.4, 50 mM NaCl) supplemented with additional reagents, as stated in the text. The experiments were performed using an in-house developed confocal microscope similar to the setup described before[27] with minor modifications as outlined below. Briefly, two laser-diodes (Coherent Obis) with emission wavelength of 532 and 637 nm were modulated in periods of 50 μs and used for confocal excitation. Alternation between both excitation wavelengths was achieved by direct modulation of the two lasers. The laser beams were combined using a dichroic mirror (T600lprx, Chroma) and then coupled into a single-mode fiber (P3-488PM-FC-2, Thorlabs) and collimated (RC12APC-P01 Thorlabs) before entering a water immersion objective (60X, NA 1.2, UPlanSAPO 60X, Olympus). The excitation spot was focused 20 µm above the interface of glass and water solution. Typical average laser powers were 30 µW at 532 nm (~30 kW/$cm^2$) and 15 µW at 637 nm (~15 kW/$cm^2$). Excitation and emission light were separated by a dichroic beam splitter (zt532/642rpc, Chroma), mounted in an inverse microscope body (IX73, Olympus). Emitted light was focused onto a 50 µm pinhole and spectrally separated (ZT640rdc, Chroma) onto two APDs (τ-spad, <50 dark-counts/sec, Excelitas Technologies) with appropriate spectral filtering (donor channel: FF01-582/75 Semrock; acceptor channel: ET700/75m Chroma). Photon arrival times were registered by a NI-Card (PCI-6601, National Instruments). Data acquisition was performed using the smfBox software[83], and initial processing was carried out using established tools. Specifically, dual-colour burst search was conducted with parameters M = 15, T = 500 µs, and L = 25 to identify fluorescence bursts. For subsequent analysis of smFRET data acquired using the smfBox platform, we utilized the FRETBursts Python package[84] Additionally, we employed the ALEX-suite for alternating-laser excitation (ALEX) data preprocessing and visualization as previously described[27]. Additional thresholding was applied to remove spurious changes in fluorescence intensity and selected for intense single-molecule bursts (total photons per burst > 250 unless otherwise mentioned). For equilibrium measurements binning the detected bursts into a 2D apparent FRET/S histogram (61 x 61 bins unless otherwise mentioned) allowed the selection of the donor and acceptor labelled molecules. The selected apparent FRET histograms (61 x 1 bins unless otherwise mentioned) were fitted using a Gaussian function. Histograms were derived from data collected from two independent protein purification procedures and of at least 3 replicates yielding more than 5000 single-molecules analysed per condition producing more than 150 Events (peak of the Gaussian function) per condition in order to fit unambiguously the Gaussian distributions. For non-equilibrium experiments, the same thresholding criteria were applied; however, Gaussian fitting was performed using the tvGMM algorithm, specifically adapted for non-equilibrium processes as described below. To initiate folding, proteins were rapidly diluted from 6 M guanidinium hydrochloride (GuHCl) into aqueous buffer. This >$10^6$-fold dilution effectively eliminated residual denaturant, allowing refolding to proceed under near-physiological conditions. smFRET measurements were started immediately after dilution, capturing apparent FRET efficiency *vs.* time trajectories. Although the optical setup closely follows the configuration previously described, targeted enhancements in optical alignment, fiber coupling efficiency, and detection pathway optimization have been implemented. These modifications result in a markedly improved photon collection efficiency, yielding approximately twice the number of detected photons per burst under otherwise identical experimental conditions. Single-molecule bursts from donor-only labelled MBP proteins were obtained by selecting the donor-only subpopulation (S > 0.9) from the 2D apparent FRET/S histogram. The total photon count per burst were normalised by its respective duration to obtain the photon count rate.

***NEXT-FRET analysis***

Apparent FRET *vs.* time datasets from folding were analyzed using the tvGMM framework, which is well-suited for handling non-equilibrium data. Prior knowledge was incorporated into the analysis to guide model convergence. Specifically, equilibrium measurements of native MBP and pre-MBP were used to extract the means (μ) and standard deviations (σ) of the corresponding Gaussian distributions. These parameters were introduced as strong priors (strength = 1.5) to anchor the native state in the analysis. All other states were left unbiased in terms of their mean positions but constrained by prior knowledge on the Gaussian widths (strength = 1), reflecting the intrinsic resolution limit (i.e. shot-noise)[68] of the microscope rather than differences in folding intermediates. The analysis was therefore guided by two key priors: (i) the expected presence of the native folded state during the folding trajectory, and (ii) the Gaussian width distribution, constrained within the experimentally observed range of 0.4–0.8, with a typical shot-noise limited width around 0.6. The initial positions of all states were randomly assigned, evenly distributed across the full range of apparent FRET values. This ensured a generic and unbiased starting point for the model, minimizing initialization bias.

### Supplementary Note 2

#### *Mathematical notation*

We summarize the glossary of symbols and the corresponding explanations relevant to the tvGMM approach in the following table.

Notation and definitions from the Methods section.

| Variable | Explanation |
|---|---|
| $T_0, T_1$ | Initial and final time of the smFRET experiment |
| $t, t_i$ | Detection time of a single molecule |
| $e, e_i$ | FRET efficiency value |
| $\mathcal{D}$ | Dataset of all FRET measurements and detection times |
| $N := \lvert\mathcal{D}\rvert$ | Total number of observed molecules |
| $p(e, t)$ | Joint probability density of FRET efficiency and detection time |
| $p(e \mid t)$ | Conditional probability density of FRET efficiency given detection time |
| $p(t)$ | Probability density of detection times |
| $Z$ | Latent variable representing the folding intermediate state |
| $K$ | Total number of distinct folding intermediate states |
| $c(t)$ | Vector of basis functions evaluated at time $t$ |
| $d$ | Dimension of the basis vector $c(t)$ |
| $p(Z = k \mid t), p_k(t)$ | Conditional probability of folding intermediate $k$ at detection time $t$ |
| $p_k(t; W)$ | Softmax-based parametrization of $p_k(t)$ |
| $W, \tilde{W}$ | Weight matrix used in the softmax function with dimension $K \times d$ |
| $p(e \mid Z = k), p_k(e)$ | Conditional probability of FRET efficiency given folding intermediate $k$ |
| $p_k(e; \mu_k, \sigma_k^2)$ | Gaussian parametrization of $p_k(e)$ |
| $\mu_k, \sigma_k^2$ | Mean and variance of the Gaussian component for state $k$ |
| $\theta, \bar{\theta}, \theta_0, \theta^{(j)}$ | Collection of all model parameters |
| $l(\theta; \mathcal{D})$ | Log-likelihood function for the dataset |
| $l_p(\theta; \mathcal{D})$ | Log-posterior incorporating prior terms |
| $\mu_{0k}, \sigma_{0k}^2$ | Prior mean and scale parameters for $\mu_k$ and $\sigma_k^2$ |
| $\lambda_\mu, \lambda_\sigma$ | Strength of the prior on $\mu_k$ and on variance $\sigma_k^2$ |
| $\gamma_{i,k}, \gamma_{i,k}(\theta)$ | Responsibility of component $k$ for observation $i$ |
| $Q(\theta \mid \bar{\theta})$ | EM lower-bound function (E-step objective) |
| $\mathcal{H}(W \mid \bar{\theta})$ | Objective function for the weight matrix in EM |
| $\epsilon$ | Stopping threshold for EM |
| $\beta_1, \beta_2, \beta_3$ | Matching coefficients |

#### *tvGMM as a generative mechanism*

The proposed tvGMM can be also seen as a generative mechanism for producing synthetic smFRET data. The relationships between the three random variables, time $T$, latent folding intermediate $Z$ and apparent FRET efficiency $E$ can be represented as a Bayesian network as shown in the left panel of the figure below. This directed graph represents the dependency structure among the three variables encoding that the apparent FRET efficiency $E$ is conditionally independent of time $T$ given the folding intermediate $Z$. The middle panel presents the evolution of folding intermediate states as parametrized by $p_k(t; W)$. Each color represents a different intermediate state, illustrating how the probabilities of each state change over a period of 500 seconds. The ground truth values for $W$ is given in the following table. Notice that only the affine parameters (e.g., the constant and the linear terms) have non-zero values. The table also reports the mathematical expressions for the basis functions for two cases: quadratic B-splines and Fourier modes. The right panel presents synthetic smFRET data generated from tvGMM where points are color-coded to match the corresponding folding intermediate from the middle panel, indicating that points of the same color are generated from the respective state.

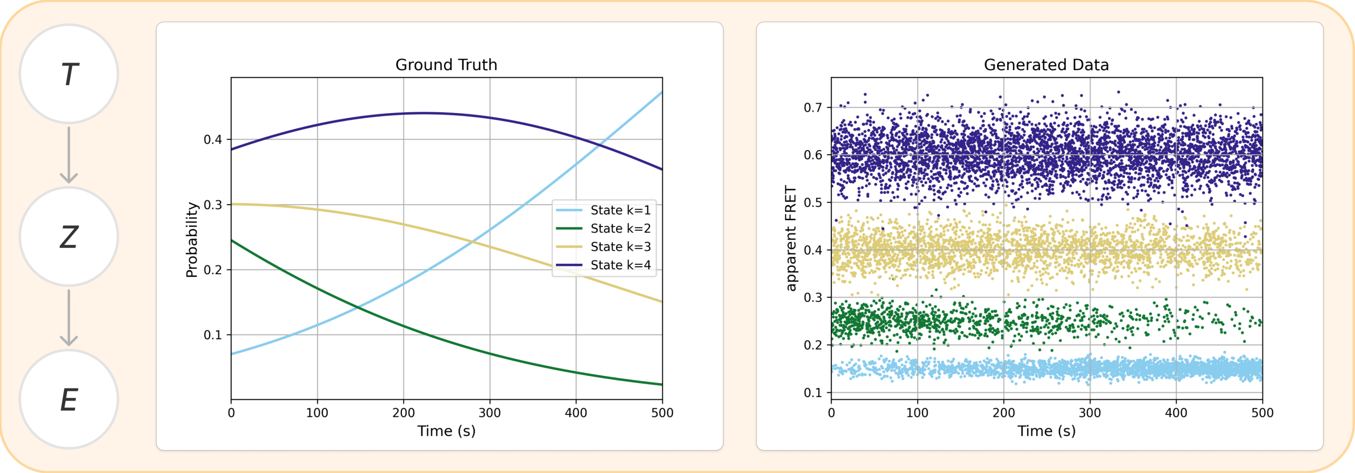


**Left**: The Bayesian network of tvGMM for the description of non-equilibrium smFRET data. **Middle**: A synthetic example with the temporal evolution of the probability for each folding intermediate state. **Right**: $N = 10^4$ synthetic smFRET data generated from tvGMM where each data point represents a FRET efficiency $e$ at a given time $t$.

Mathematical formulas for the basis functions and ground truth values for $\theta$ that generate the smFRET measurements of the above figure. For notational compactness, we change the time variable to $s = (t - T_0)/(T_1 - T_0)$.

| | $c_1(s)$ | $c_2(s)$ | $c_3(s)$ | $c_4(s)$ | $c_5(s)$ |
|---|---|---|---|---|---|
| Affine + B-Splines | 1 | $s$ | $(1-s)^2$ | $2s(1-s)$ | $s^2$ |
| Affine + Fourier | 1 | $s$ | $\cos(2\pi s)$ | $\sin(2\pi s)$ | $\cos(4\pi s)$ |

| **State** | $w_1$ | $w_2$ | $w_3$ | $w_4$ | $w_5$ | $\mu_k$ | $\sigma_k^2$ |
|---|---|---|---|---|---|---|---|
| $k = 1$ | 0.00 | 0.00 | 0.00 | 0.00 | 0.00 | 0.15 | 0.01 |
| $k = 2$ | -1.25 | 8.51 | 0.00 | 0.00 | 0.00 | 0.25 | 0.02 |
| $k = 3$ | -1.45 | 5.19 | 0.00 | 0.00 | 0.00 | 0.40 | 0.03 |
| $k = 4$ | -1.70 | 3.98 | 0.00 | 0.00 | 0.00 | 0.60 | 0.04 |

***Quantitative demonstrations on synthetic data***

We performed a series of synthetic experiments to evaluate the parameter estimation accuracy of the tvGMM framework. Synthetic smFRET datasets were generated using the ground truth parameters listed in the above table, with sample sizes ranging from 1k to 100k. The left panel of the figure below illustrates the estimated state occupancy probabilities alongside the ground truth for the case of 10k samples, using affine functions combined with B-splines as the basis. The close alignment between the estimated and true curves demonstrates the model's ability to recover the underlying state progression over time and indicates the accuracy of the estimated parameter matrix $W$. The right panel presents a comparison between two basis function families: B-splines (red dashed line) and Fourier modes (orange dashed line). Both representations demonstrate a reduction in Mean Absolute Error (MAE) as the number of single molecules increases. However, B-splines consistently outperform Fourier modes, achieving lower MAE across all sample sizes making it the default choice for our basis vector function $c(t)$. It is also worth noting that the rate of error reduction matches the expected theoretical convergence behavior, shown by the black dashed line.

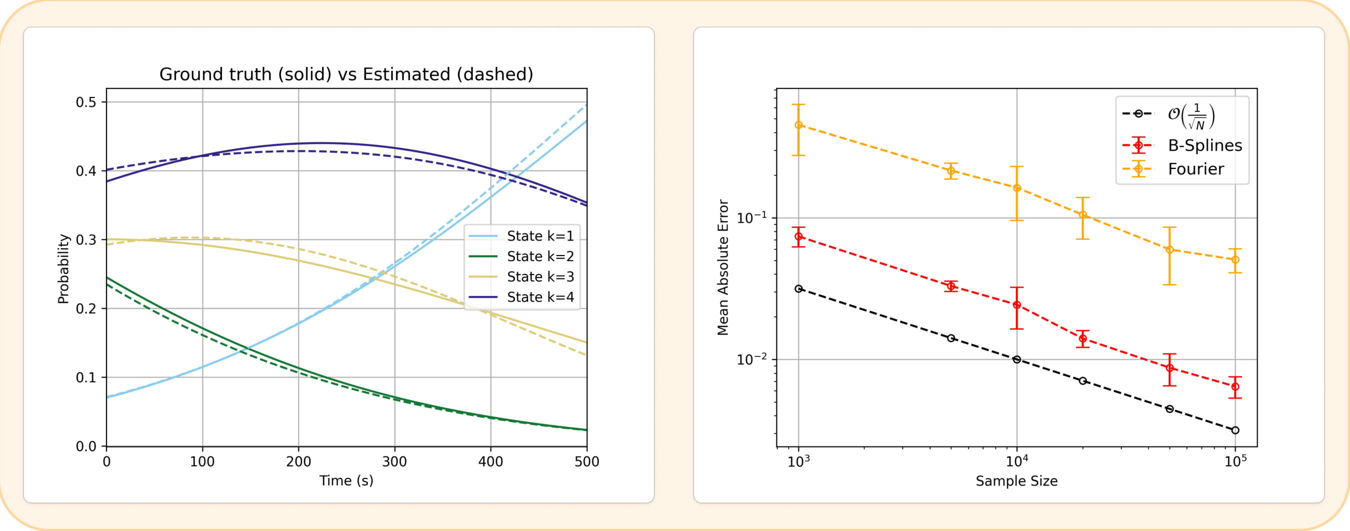


**Left:** Comparison of the estimated folding intermediate probabilities with the ground truth probabilities for 10k samples. **Right:** Mean Absolute Error (MAE) between the true and estimated probabilities with respect to the sample size.

We next investigate the influence of prior knowledge on the estimation of the Gaussian component parameters. In particular, we explore how varying the strength of the priors affects the resulting estimates. As the prior strength parameters $\lambda_\mu$ and $\lambda_\sigma$ increase, the corresponding prior distributions become increasingly concentrated around their means. This reflects an increasing level of confidence in the prior, resulting in lower variance in the parameter estimates. To study this effect systematically, we define the prior strength as power functions of the sample size, $N$,

$$\lambda_\mu = N^p \quad \text{and} \quad \lambda_\sigma = N^p$$

with the exponent $p$ determining how strongly the prior scales with data availability.

We assessed the impact of prior knowledge under two experimental design factors: sample size and the presence of noise contamination in smFRET data. Synthetic datasets were generated with varying levels of background noise (0%, 10%, and 20%), sampled from the uniform distribution $\mathcal{U}[0.1, 0.7]$. The left column of the figure below illustrates simulated smFRET data over time, color-coded by folding intermediate state, across different noise levels with noisy samples shown in red. The middle and right columns summarize the MAE in estimating means and variances, computed over 50 repetitions for each combination of sample size ($N = 50, 100, 500, 1000$) and prior strength exponent ($p = 0, 0.5, 1$). The results demonstrate that stronger priors (i.e., larger $p$) consistently reduce estimation error and its variance, particularly in low-data and high-noise conditions. Importantly, while increasing sample size consistently improves estimation accuracy, incorporating appropriately scaled priors substantially mitigates the adverse effects of noise, enhancing the stability of inference in high-uncertainty regimes. Across both panels, we observe a monotonic relationship between prior strength and estimation accuracy: stronger priors (larger exponents) result in consistently lower estimation errors. This trend holds across all noise levels and sample sizes, with higher-power priors outperforming weaker ones. These results empirically validate the benefit of using informed priors that scale with sample size, particularly when data are limited and confidence in prior knowledge is high.

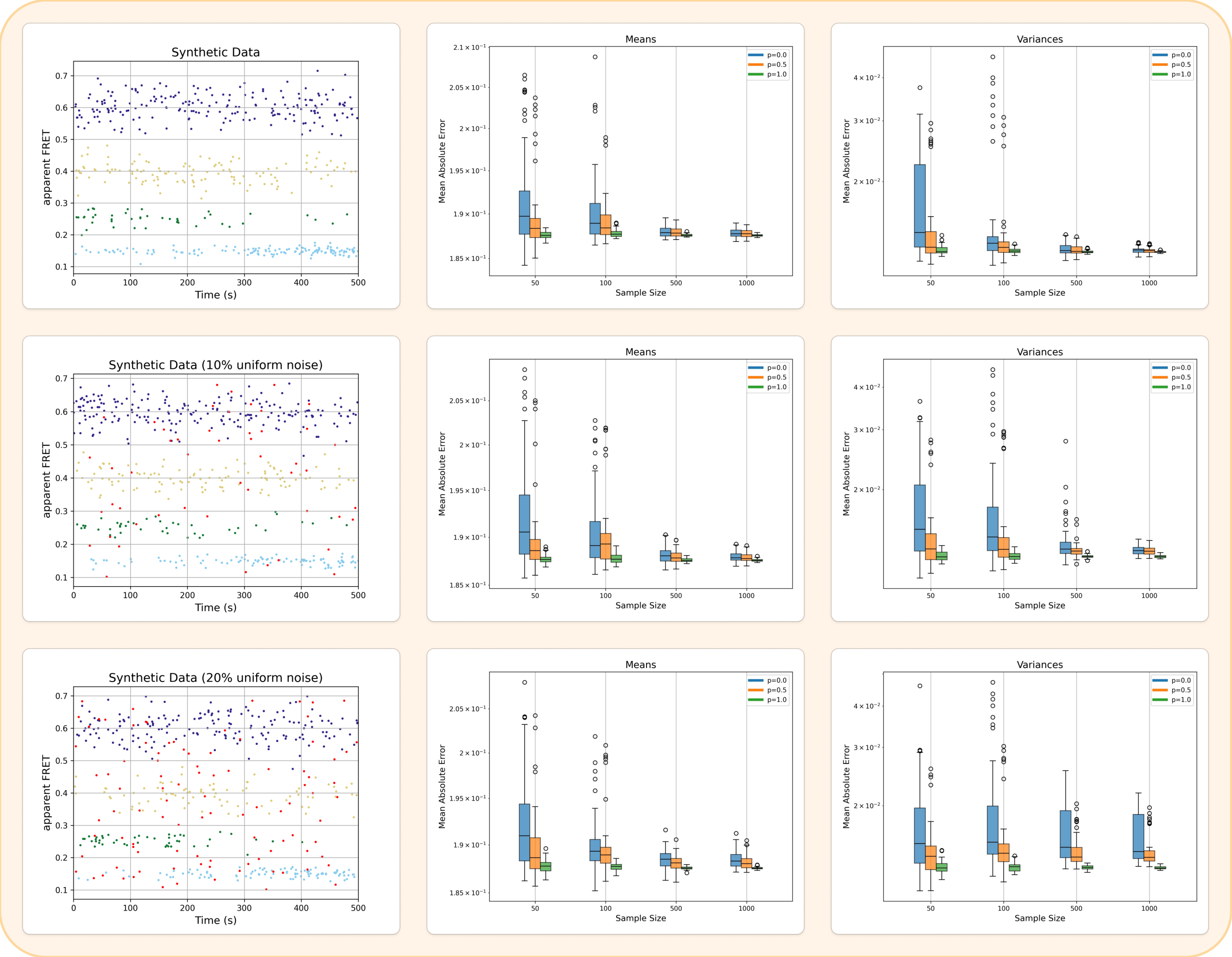


**Left:** Generated smFRET data with three different levels of uniform noise. **Middle:** MAE between the true and estimated mean vector across varying sample sizes and prior strength power $p$. **Right:** The corresponding MAEs for the estimated variance vector.

***Selecting the number of folding intermediate states***

To assess model selection performance, we generated synthetic datasets of varying sample sizes ($N$ = 1k, 5k, 10k & 20k) using a ground truth model with $K = 4$ components. For each dataset, we evaluated BIC values across models with $K = 2$ to 6. In all cases, the minimum BIC was observed at $K = 4$, successfully recovering the true number of components, as shown in the table below. These results were obtained without incorporating prior knowledge for the estimation, however, the same selection strategy applies when priors are used. Similar results are obtained when 20% uniform noise is added to the dataset showing the robustness of BIC against background noise.

BIC values across different sample sizes and numbers of states. Lower values indicate better model fit.

| Sample Size | $K = 2$ | $K = 3$ | $K = 4$ | $K = 5$ | $K = 6$ |
|---|---|---|---|---|---|
| 1k | -1350 | -1744 | **-1956** | -1908 | -1872 |
| 5k | -6932 | -9086 | **-10317** | -10258 | -10213 |
| 10k | -14015 | -18178 | **-20671** | -20613 | -20561 |
| 20k | -28243 | -36683 | **-41815** | -41577 | -41679 |

***Detailed derivation of the formulas in the EM algorithm***

By introducing an auxiliary distribution $q_k(t)$ and applying Jensen's inequality to the tvGMM's log-likelihood defined in Eq. (4) of the Methods, we obtain the following lower bound:

$$
\begin{aligned}
l(\theta; \mathcal{D}) &= \sum_{i=1}^{N} \log \left( \sum_{k=1}^{K} p_k(t; W) p_k(e_i; \mu_k, \sigma_k^2) \right) = \sum_{i=1}^{N} \log \left( \sum_{k=1}^{K} \frac{p_k(t; W) p_k(e_i; \mu_k, \sigma_k^2)}{q_k(t)} \right) \\
&\geq \sum_{i=1}^{N} \sum_{k=1}^{K} q_k(t) \log \frac{p_k(t; W) p_k(e_i; \mu_k, \sigma_k^2)}{q_k(t)} = \sum_{i=1}^{N} \sum_{k=1}^{K} q_k(t) \log p_k(t; W) p_k(e_i; \mu_k, \sigma_k^2) + const.
\end{aligned}
$$

The inequality holds for any choice of $q_k(t)$, but becomes an equality when $q_k(t) = \gamma_{i,k}(\theta)$, the posterior responsibility under the current parameter estimate.

We now derive the update equations for the parameters of the Gaussian components. By differentiating Eq. (10) from the Methods section with respect to $\mu_k$ and equating the derivative to zero, we obtain the following closed-form solution:

$$
\begin{aligned}
&\frac{\partial}{\partial \mu_k} \left( \sum_{i=1}^{N} \sum_{k=1}^{K} \gamma_{i,k} \log \left( p_k(t_i; W) p_k(e_i; \mu_k, \sigma_k^2) \right) - \frac{\lambda_\mu}{2} \sum_{k=1}^{K} (\mu_k - \mu_{0k})^2 - \frac{\lambda_\sigma}{2} \sum_{k=1}^{K} \left( \log \sigma_k^2 + \frac{\sigma_{0k}^2}{\sigma_k^2} \right) \right) = 0 \\
&\Rightarrow \sum_{i=1}^{N} \frac{\gamma_{i,k}(e_i - \mu_k)}{\sigma_k^2} - \lambda_\mu (\mu_k - \mu_{0k}) = 0 \\
&\Rightarrow \hat{\mu}_k = \frac{\sum_{i=1}^{N} \gamma_{i,k} e_i + \lambda_\mu \sigma_k^2 \mu_{0k}}{\sum_{i=1}^{N} \gamma_{i,k} + \lambda_\mu \sigma_k^2}
\end{aligned}
$$

The variance term $\sigma_k^2$ that appears in the update equation can be absorbed into $\lambda_\mu$. This adjustment enhances numerical stability—particularly in cases where small values of $\sigma_k^2$ would otherwise diminish the influence of the prior, resulting in less robust estimates. Incorporating this consideration, the estimate for the mean of the $k$-th component becomes:

$$
\hat{\mu}_k = \frac{\sum_{i=1}^{N} \sum_{k=1}^{K} \gamma_{i,k} e_i + \lambda_\mu \mu_{0k}}{\sum_{i=1}^{N} \sum_{k=1}^{K} \gamma_{i,k} + \lambda_\mu}
$$

Likewise, differentiating Eq. (10) from the Methods section with respect to $\sigma_k^2$ and equating the result to zero yields the following closed-form expression:

$$
\begin{aligned}
&\frac{\partial}{\partial \sigma_k^2}\left(\sum_{i=1}^{N}\sum_{k=1}^{K}\gamma_{i,k}\log\left(p_k(t_i;W)p_k(e_i;\mu_k,\sigma_k^2)\right)-\frac{\lambda_\mu}{2}\sum_{k=1}^{K}(\mu_k-\mu_{0k})^2-\frac{\lambda_\sigma}{2}\sum_{k=1}^{K}\left(\log\sigma_k^2+\frac{\sigma_{0k}^2}{\sigma_k^2}\right)\right)=0\\
&\Rightarrow\sum_{i=1}^{N}\gamma_{i,k}\left(\frac{(e_i-\mu_k)^2}{2\sigma_k^4}-\frac{1}{2\sigma_k^2}\right)-\frac{\lambda_\sigma}{2}\left(\frac{1}{\sigma_k^2}-\frac{\sigma_{0k}^2}{\sigma_k^4}\right)=0\\
&\Rightarrow\sigma_k^2\left(\sum_{i=1}^{N}\gamma_{i,k}+\lambda_\sigma\right)=\sum_{i=1}^{N}\gamma_{i,k}(e_i-\mu_k)^2+\lambda_\sigma\sigma_{0k}^2\\
&\qquad\qquad\qquad\qquad\Rightarrow\hat{\sigma}_k^2=\frac{\sum_{i=1}^{N}\gamma_{i,k}(e_i-\mu_k)^2+\lambda_\sigma\sigma_{0k}^2}{\sum_{i=1}^{N}\gamma_{i,k}+\lambda_\sigma}\,.
\end{aligned}
$$

**Supplementary Figures**

Supplementary Figure 1. Enhanced resolution of MBP folding intermediated by NEXT-FRET compared to traditional histogram-based analysis

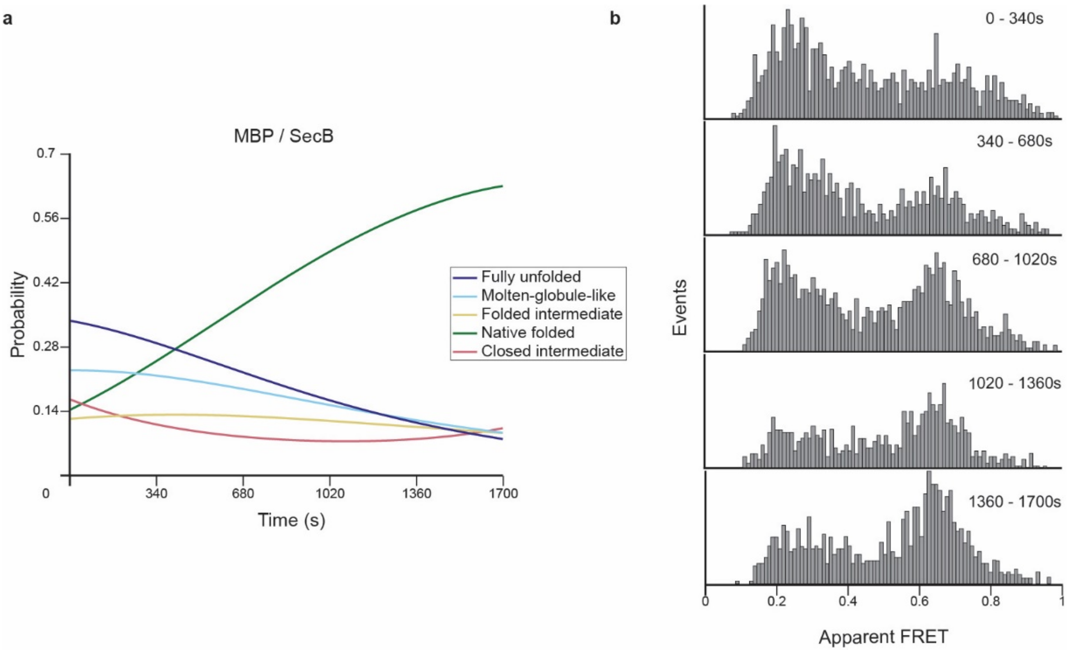


**a**, Time-resolved smFRET analysis of MBP folding in the presence of SecB, using NEXT-FRET. This method infers intermediate state probabilities directly from unbinned FRET *vs.* time data, enabling high-resolution tracking of folding trajectories and detection of low-abundance intermediates with minimal bias. **b**, The same dataset analyzed using conventional equilibrium-based Gaussian fitting approach. Events were binned into 340s intervals to enable analysis, but this temporal averaging masks dynamic transitions, blurs state separation, and obscures minor populations. These results demonstrate the advantages of NEXT-FRET over traditional approaches for resolving folding intermediates in complex, dynamically evolving single-molecule trajectories; particularly in the presence of chaperones like SecB.

Supplementary Figure 2. Chaperones with negligible impact on MBP and pre-MBP folding dynamics.

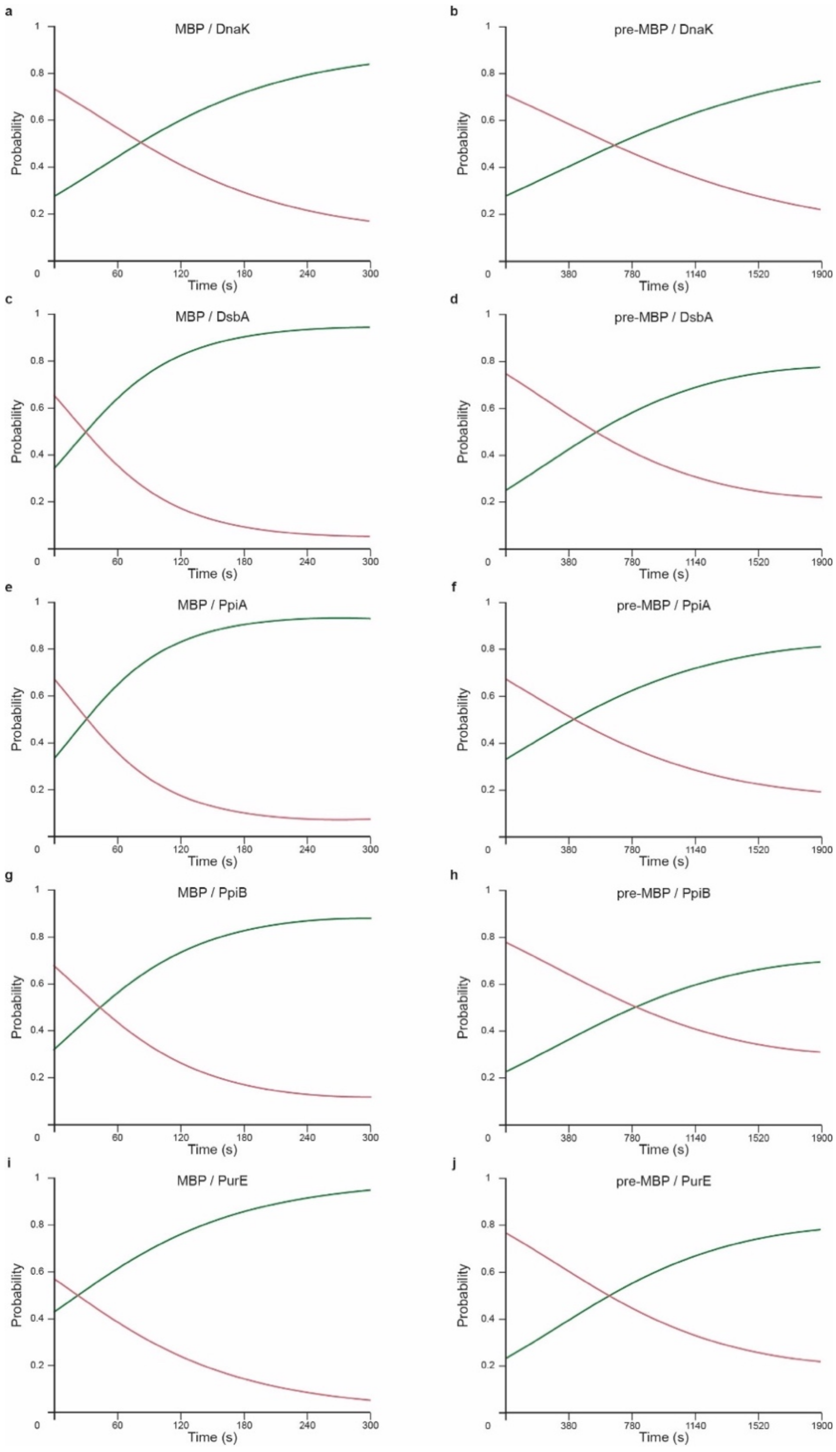


Time-dependent smFRET trajectory analysis reveal that DnaK (**a–b**), DsbA (**c–d**), PpiA (**e–f**), PpiB (**g–h**), and PurE (**i–j**) exhibit minimal or no impact on the folding behavior of MBP (left panels: a, c, e, g, i) or pre-MBP (right-panels: b, d, f, h, j). State occupancies over time remain comparable to unassisted folding, with no notable shifts in intermediate or native-state populations. These results validate the specificity of chaperone modulation observed with SecB, SecA, TF and Skp, and validate the robustness of our analytical approach across a wide range of experimental conditions.

### Supplementary Tables

Supplementary Table 1. Model selection across all primary datasets using Bayesian Information Criterion (BIC).

| Folding states | 2 | 3 | 4 | 5 | 6 |
|---|---|---|---|---|---|
| MBP | **-1246.56** | -3721.37 | -3684.32 | - | - |
| pre-MBP | **-2634.22** | -4983.89 | -5025.34 | -5071.52 | - |
| MBP / Tf | 56.88 | **-3472.23** | -3998.43 | -3969.81 | - |
| pre-MBP / Tf | - | -2187.89 | **-2565.99** | -2551.23 | - |
| MBP / Skp | - | -2980.94 | **-3659.83** | -3520.65 | - |
| MBP / secA | - | -1460.18 | **-1540.84** | -1473.71 | - |
| pre-MBP / secA | - | -103.42 | **-885.19** | -909.83 | -948.86 |
| MBP / secB | - | - | -4356.80 | **-4427.80** | -4335.98 |
| pre-MBP / secB | - | - | -2564.06 | **-3050.69** | -2992.50 |

BIC values were calculated for each dataset across models with 2–6 folding states. Cells with bold font indicate the final number of states selected for downstream NEXT-FRET analysis. Red font cells highlight the BIC-optimal state number when it differs from the selected model. In such cases, additional states—although statistically favored—were attributed to noise coming from random coincident fluorophore events or photobleaching[68]. Final model state selections reflect a balance between statistical optimality and physical plausibility, ensuring robust and biologically meaningful folding intermediate assignments.

Supplementary Table 2. Apparent FRET efficiencies and standard deviations for all identified folding states.

| Folding states | Standard Deviation Apparent FRET |
|---|---|
| Fully Unfolded | 0.23 ± 0.05 |
| Molten-globule-like | 0.32 ± 0.08 |
| Folded Intermediate | 0.47 ± 0.05 |
| Native Folded | 0.64 ± 0.02 |
| Closed Intermediate | 0.83 ± 0.04 |
